# Complex biophysical changes and reduced neuronal firing in an *SCN8A* variant associated with developmental delay and epilepsy

**DOI:** 10.1101/2023.12.04.569940

**Authors:** Shir Quinn, Nan Zhang, Timothy A. Fenton, Marina Brusel, Preethi Muruganandam, Yoav Peleg, Moshe Giladi, Yoni Haitin, Holger Lerche, Haim Bassan, Yuanyuan Liu, Roy Ben-Shalom, Moran Rubinstein

## Abstract

**Background:** Mutations in the *SCN8A* gene, encoding the voltage-gated sodium channel Na_V_1.6, lead to various neurodevelopmental disorders. The *SCN8A* p.(Gly1625Arg) mutation (Na_V_1.6^G1625R^) was identified in a patient diagnosed with developmental epileptic encephalopathy (DEE), presenting with moderate epilepsy and severe developmental delay.

**Methods:** We performed biophysical and neurophysiological characterizations of Na_V_1.6^G1625R^ in Neuro-2a cells and cultured hippocampal neurons, followed by computational modeling to determine the impact of its heterozygous expression on cortical neuron function.

**Findings:** Voltage-clamp analyses of Na_V_1.6^G1625R^ demonstrated a heterogeneous mixture of gain-and loss-of-function properties, including reduced current amplitudes, a marked increase in the time constant of fast voltage-dependent inactivation and a depolarizing shift in the voltage dependence of inactivation. Recordings in transfected cultured neurons showed that these intricate biophysical properties had a minor effect on neuronal excitability when firing relayed on both endogenous and transfected Na_V_ channels. Conversely, there was a marked reduction in the number of action potentials when firing was driven by the transfected mutant Na_V_1.6 channels. Computational modeling of mature cortical neurons further revealed a mild reduction in neuronal firing when mimicking the patients’ heterozygous Na_V_1.6^G1625R^ expression. Structural modeling of Na_V_1.6^G1625R^ and a double-mutant cycle analysis suggested the possible formation of pathophysiologically-relevant cation-π interaction between R1625 and F1588, affecting the voltage dependence of inactivation.

**Interpretation:** Our analyses demonstrate a complex combination of gain and loss-of-function changes resulting in an overall mild reduction in neuronal firing, related to a perturbed interaction network within the voltage sensor domain.

**Funding:** ISF, DFG, BMBF, The Hartwell Foundation, ICRF, ISCA

**Research in context:** *Evidence before this study:* Mutations in the *SCN8A* gene, encoding the voltage-gated sodium channel Na_V_1.6, result in multiple neurodevelopmental syndromes ranging from benign epilepsy to developmental delay without epilepsy or developmental epileptic encephalopathy (DEE). Recent studies established that most DEE-causing *SCN8A* mutations result in a gain of function effect. However, several *SCN8A* mutations that diverge from this pattern were described.

*Added value of this study:* We performed a multi-tiered study of the *SCN8A* p.(Gly1625Arg) variant (Na_V_1.6^G1625R^), identified in a patient with atypical DEE presentation, featuring moderate epilepsy that is well controlled by the sodium channel blocker Oxcarbazepine, along with profound stagnated developmental delay. This variant is positioned within the S4 segment of domain IV, a critical region for Na_V_1.6 function, where pathogenic variants were shown to cause either a loss or gain of channel function, but often with mixed biophysical alterations. Our biophysical characterization of Na_V_1.6^G1625R^ in Neuro-2a cells demonstrated complex gain-and loss-of-function properties, cumulating to reduced firing in cultured hippocampal neurons and computational modeling of mature cortical neurons, demonstrating an overall loss-of-function effect.

*Implications of all the available evidence:* Our results indicate the necessity for combined biophysical and neuronal characterization of individual *SCN8A* variants, especially those presenting with complex biophysical changes or atypical clinical presentation. Moreover, while sodium channel blockers are the recommended treatment for *SCN8A* variants associated with gain-of-function, additional considerations may be needed for DEE-causing variants that are associated with mild loss-of-function.

## Introduction

Voltage[gated sodium channels (Na_V_s) are key regulators for neuronal excitability, mediating the generation and propagation of action potentials. In line with their critical contribution to neuronal firing, pathogenic variants in the genes coding for brain-expressed Na_V_ channels (*SCN1A*, *SCN2A*, *SCN3A,* and *SCN8A*) are strongly associated with neurological disorders [1].

The *SCN8A* gene encodes for Na_V_1.6. These channels comprise of four homologous domains (DI-DIV), each containing six transmembrane segments (S1-S6). Within each domain, the transmembrane segments S1-S4 form the voltage-sensor subdomain (VSD), while S5-S6 constitute the pore-forming subdomain [1]. Pathogenic *SCN8A* variants have been implicated in a range of neurodevelopmental disorders, including autism spectrum disorder, intellectual disability, benign familial infantile epilepsy (BFIE), and developmental and epileptic encephalopathy (DEE) [2,3]. Most mutations occurred de novo, and over 90% are missense mutations [3]. Functional analyses that addressed *SCN8A*-related genotype-phenotypes relations demonstrated that gain of function (GoF) variants are associated with mild or intermediated focal epilepsy and DEE, whereas loss of function (LoF) variants are related to generalized epilepsy and neurodevelopmental disorders without epilepsy [3–5].

Johannesen et al. (2022) reported on a cohort of 392 individuals harboring different *SCN8A* mutations, most of which are missense mutations. Functional analysis was performed on multiple mutations, indicating that *SCN8A* variants associated with DEE show a propensity for GoF properties due to shifts in the voltage dependency of activation or inactivation, a slower rate of inactivation, and an increased persistent current (Supplementary Table 1)[3,5,14–19,6–13].

The recurrent *SCN8A* missense c.4873G>A mutation, converting glycine 1625 to arginine in the S4 segment of the domain IV (Na_V_1.6^G1625R^), has been independently identified in three patients with DEE [3,20,21]. This mutation was classified as pathogenic, predicted to cause GoF (67%) [22–24], but was not functionally characterized.

Here, we performed an in-depth analysis demonstrating biophysical changes, altered neuronal firing, and the involvement of electrostatic molecular interaction with the voltage sensor domain of domain IV that is involved in these functional ramifications. These data indicate the complex functional consequences inflicted by DEE-associated *SCN8A* mutations.

## Methods

### DNA plasmids

The TTX-insensitive human Na_V_1.6 was used before [3,5]. G1625R and F1588A mutations were introduced by overlap extension-PCR, followed by restriction enzyme digestion (*XhoI* and *Hind*III) and ligation. For G1625R mutation SCN8A_G1625R_F and SCN8A_seq6F primers and SCN8A_G1625R_F and SCN8A_R primers were used (Table 1) in two separate PCR reactions. The PCR products were purified (Qiagen PCR purification kit) and assembled using the flanking primers, SCN8A_seq6F and SCN8A_R. The combined PCR product was digested with *XhoI* and *Hind*III and ligated to pCDNA6-SCN8A-WT vector digested with the same restriction enzymes. The introduction of F1588A mutation was essentially identical, except that SCN8A_F1588A_F and SCN8A_F1588A_R primers were used (Table 1). For generation of clone harboring two mutations (F1588A and G1625R), pCDNA6-SCN8A-G1625R was used as a temple for the introduction of F1588A mutation, as described above for the single mutation.

**Table 1.**
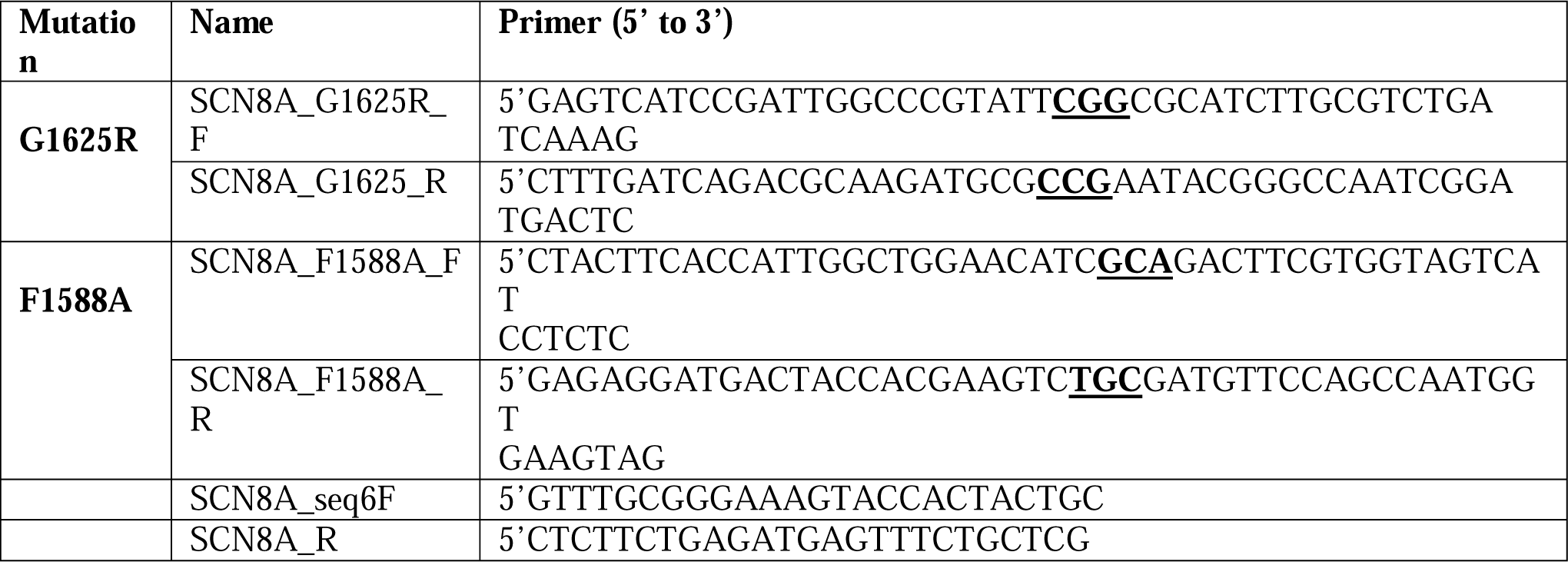
Primers used for site-directed mutagenesis.

All DNA manipulations were performed in *E. coli* CopyCutter (Lucigen, C400CH10). The sequence was verified using sequencing of the full coding sequence.

### Transfection and expression in Neuro-2a cells

The murine neuroblastoma cell line Neuro 2a, was cultured in Dulbecco’s Modified Eagle Medium supplemented with 10% fetal bovine serum, L-glutamine (2 mM), 10 units/mL penicillin and 10μg/mL streptomycin (Biological Industries, Beit-Haemek, Israel), and grown at 37 °C with 5% CO_2_. The cells were transfected using X-tremeGENE 9 (Sigma-Aldrich) with 3.2 mg Na_V_1.6 (Na_V_1.6^WT^, Na_V_1.6^G1625R^, Na_V_1.6^F1588A^, Na_V_1.6^F1588A,G1626R^) together with 0.32 mg of human β1 (pCLH-hb2-CD8) and 0.32 mg β2 (pCLH-hb1-EGFP) subunits. Electrophysiological recordings were performed 48 h after transfection from cells expressing all three subunits, which were recognized by (i) a TTX-resistant Na^+^ current (α-subunit); (ii) anti-CD8 antibody-coated microbeads on the cell surface (β1-subunit); and (iii) a green fluorescence (β2-subunit) as done before [5].

### Neuro-2a cells voltage clamp recordings

Whole-cell voltage-clamp recordings were performed as described before, but with some modifications [25]. The pipette solutions contained: 140 mM CsF, 10 mM NaCl, 5 mM EGTA, 12 mM HEPES, 10 mM glucose, adjusted to pH 7.3 with CsOH. The external solution contained: 140 mM NaCl, 5.4 mM CsCl, 1.8 mM CaCl_2_, 1 mM MgCl_2_, 10 mM HEPES, adjusted to pH 7.4 with NaOH. Chemicals were purchased from Sigma-Aldrich or Fisher Chemical. 500 mM TTX (Alomone labs, Cat# T-550) was added to the bath solution to block endogenous Na_V_ channels.

Recordings were made using Sutter IPA amplifier (Sutter Instrument, Novato) sampled at 50 kHz and filtered 10 kHz. The glass pipettes had resistances of 3–5 MΩ. Capacitive currents were minimized using the amplifier circuitry. We routinely used 75–80% series resistance compensation. The remaining linear capacity and leakage currents were eliminated by P/4 subtraction.

To measure the voltage dependence of activation, cells were held at -120 mV and depolarized from -80 mV to +30 mV with 10 mV voltage increments. V_1/2_ was derived from the Boltzmann function [25]. The time constant for channel inactivation was calculated by fitting a monoexponential function. The voltage dependence of inactivation was measured from a holding potential of -120 mV, cells were depolarized for 100 ms from -140 mV to 0 mV with 10 mV increments and held at -10mV for 50 ms after each voltage step before returning to holding potential. V_1/2_ was derived from the Boltzmann function. To measure ramp currents, the voltage was changed from a holding of -120 mV to -60 mV, increasing gradually to 0mV over 300 ms. For the analyses, ramp currents were normalized to peak currents. The recovery from inactivation was measured from a holding of -120 mV, with a pretest pulse to -10 mV that lasted for 100 ms, followed by a recovery time ranging from 1 ms to 20 ms, in 1 ms increments at -120 mV and a test pulse at -10 mV. Repeated depolarizations (nine) were measured from a holding potential -70 mV, followed by 10 ms depolarizations to -10 mV in 10 ms intervals. Double mutant cycle analysis was done as described before [26].

### Cultured hippocampal neurons

Animal protocols for primary cell culture were approved by the local Animal Care and Use Committee (Regierungspraesidium Tuebingen, Tuebingen, Germany). Mouse hippocampal neuronal cultures were obtained and transfected as previously described [5]. Standard whole-cell patch clamp and current clamp experiments were performed from the transfected neurons using an Axopatch 200B amplifier, a Digidata 1440A digitizer and Clampex 10.2 data acquisition software at a room temperature either in absence or presence of 500 nM tetrodotoxin (TTX). Cells visualized with a Leica DM IL LED microscope were held at -70 mV. For current-clamp recordings, signals were low-pass filtered at 10 kHz and sampled at 100 kHz. Borosilicate glass pipettes had final resistances of 3 – 5 MΩ when filled with the pipette solution containing (in mM): 5 KCl, 4 ATP-Mg, 10 phosphocreatine, 0·3 GTP-Na, 10 HEPES, 125 K-gluconate, 2 MgCl_2_ and 10 EGTA, pH 7.2 with KOH, osmolarity 290 mOsm/kg with mannitol. The bath solution contained (in mM): 125 NaCl, 25 NaHCO_3_, 2.5 KCl, 1 MgCl_2_, 2 CaCl_2_, 1.25 NaH_2_PO_4_, 5 HEPES, 10 Glucose, pH 7.4 with NaOH, osmolarity 305 mOsm/kg with mannitol. The liquid junction potential was not corrected.

### Computational modeling

Simulations were performed using a compartmental model of a layer 5 pyramidal neuron initially described before [27]. This model is based on a pyramidal cell within the NEURON environment [28], incorporating morphological parameters and channel distributions in line with those delineated by Ramaswamy and Markram, (2015), and electrophysiological properties as reported before [27,28,30]. In this iteration, Na_V_1.6 channels in the distal AIS were modeled using a multi-state Hidden Markov Model (HMM), a departure from the Hodgkin-Huxley (HH) model previously employed. Na_V_1.2 channels were represented by an eight-state model [31].

The optimization of parameters for the compartmental model containing HMM Na_V_1.6 channels was conducted using an evolutionary algorithm facilitated by BluePyOpt [32], which was adapted to the computational infrastructure of the National Energy Research Computing Center [33]. This process ensured the functional equivalence of the HMM model parameters to those of the established HH model before incorporating any Na_V_1.6 mutations. After validation that the HMM model parameters emulated the physiological behaviors observed in HH models, different ratios of the Na_V_1.6^G1625R^ mutation were introduced across various neuronal compartments, including nodes of Ranvier, distal AIS, and somatodendritic regions. The resulting impact on neuronal function was evaluated against the newly established baseline activity. To model the transfection of Na_V_1.6 channels in cultured neurons, we increased the Na_V_1.6 conductance density by 20% throughout the model neuron for either Na_V_1.6^WT^ or Na_V_1.6^G1625R^ to the axon initial segment (AIS), while maintaining the endogenous expression of Na_V_1.2 and Na_V_1.6 channels constant.

### Statistics

All data are shown as mean ± standard error of the mean (SEM), and in most cases, also the individual data points. Statistical analysis utilized students’ t-test or Mann Whitney when comparing two groups. One-way ANOVA followed by Holm-Sidek multiple comparisons, or the nonparametric Kruskal-Wallis test, followed by Dunn’s multiple comparisons tests, were used when comparing more than two groups as indicated in the figure legend. The number of samples in each figure is indicated in the legend.

### Ethics statement

All analyses were performed as part of clinical care reported according to IRB approval, and consent was obtained from the parents. Animal protocols for primary cell culture were approved by the local Animal Care and Use Committee (Regierungspraesidium Tuebingen, Tuebingen, Germany).

### Role of funding source

The funders had no role in study design, data collection, data analysis or interpretation, or the decision to prepare or publish the manuscript.

## Results

### Clinical description

The missense *SCN8A* G1625R mutation was identified in a child diagnosed with DEE, presented as moderate epilepsy with severe developmental delay. The patient was born with an *in vitro* fertilization twin pregnancy at 37 weeks gestation with a birth weight of 2018 gr. and an Apgar score of 9/10. Global Hypotonia was detected since birth, and soon after, severe global developmental delay was diagnosed. Developmental quotient was < 40 at 12 months. He then presented extrapyramidal movements (chorea and dystonia) diagnosed as dyskinetic and hypotonic quadriplegic cerebral palsy with a gross motor function classification level (GMFCS) of 4-5. Magnetic resonance imaging (MRI) at 12 months showed delayed myelination, and at 28 months, additional mild white matter changes (Fig. 1A). Seizure onset was at 23 months, with a 15-second left focal clonic seizure. Additionally, his parents reported brief intermittent episodes of starring, eye deviations, and clonic unilateral or bilateral leg movement. Electroencephalography (EEG) showed non-frequent bilateral centro-parietal interictal epileptic activity, while no ictal activity was captured.

**Fig 1.**
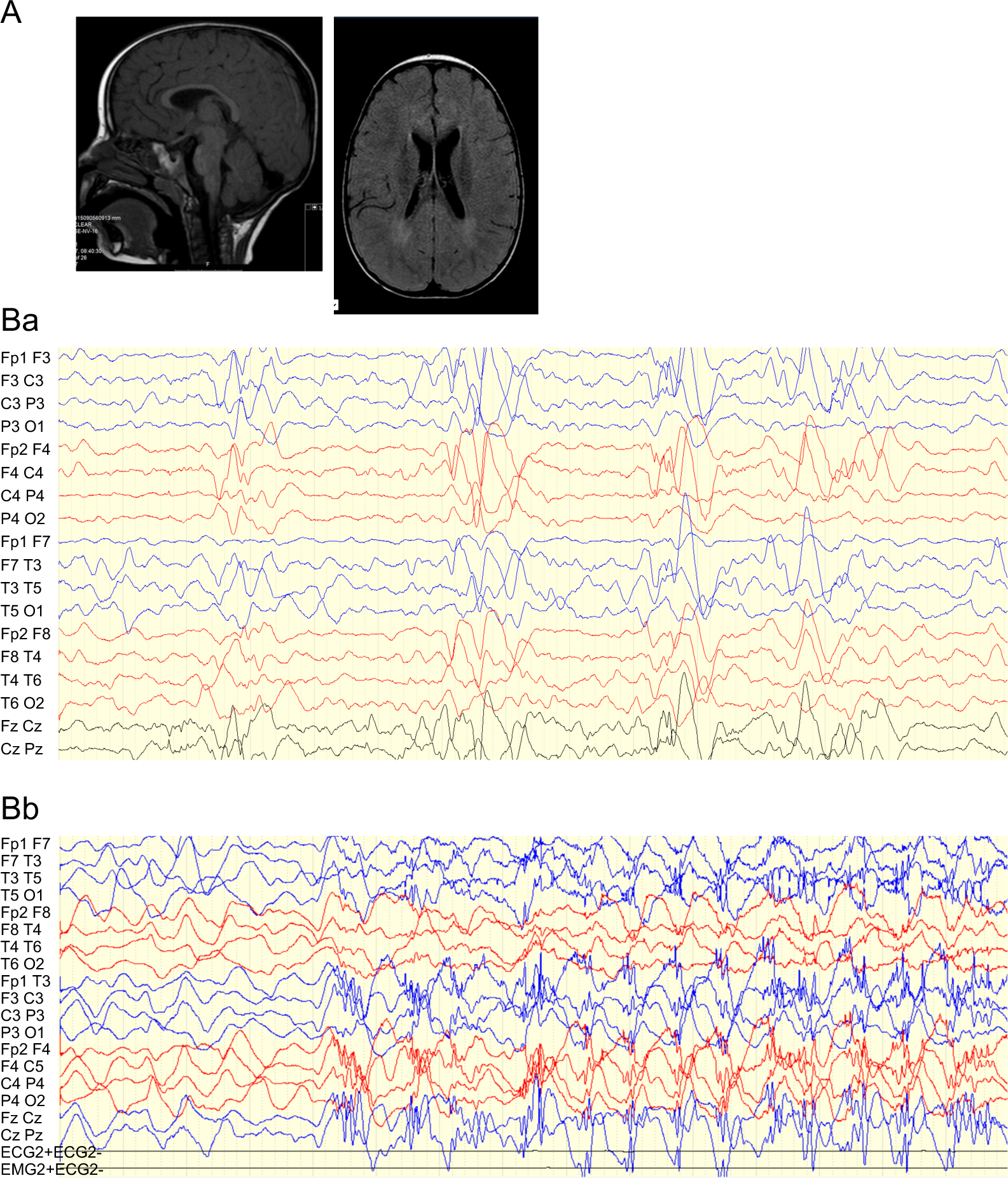
MRI and EEG features of a patient with the SCN8A G1625R mutation. **(A)** MRI scans showed mild thinning of the corpus callosum and delayed myelination. **(Ba)** EEG at three years of age showing interictal frontocentral epileptiform activity **(Bb)** EEG at six years of age, demonstrating nocturnal electroclinical tonic events with associated bilateral frontocentral polyspike activity.

The sodium channel blocker carbamazepine was initiated and was later switched to oxcarbazepine due to the inability to reach appropriate blood levels. During the following year, the parents reported a decrease in the frequency of staring / eye deviation seizures and only an additional one brief focal seizure. Video EEG (VEEG) at 3 years old demonstrated interictal frontocentral epileptiform activity without electrographic or electroclinical seizures (Fig. 1Ba).

Parallel to seizure onset, his developmental achievements were modest, including mild improvement in eye contact and ability to sit better, but overall, his development was arrested in all domains. Over time, the frequency of seizures gradually and significantly decreased, and at the age of six years, Oxcarbazepine was discontinued, but soon after, starring episodes and tonic seizures were reported. VEEG captured several nocturnal electroclinical tonic events with associated bilateral frontocentral polyspike activity (Fig. 1Bb), and interictal semi-rhythmic sharp wave activity. Oxcarbazepine was reinitiated with a significant decrease in seizure frequency.

Towards six years of age, he developed left Achilles tightness, and he transiently responded to gastrocnemius botulinum toxin injection while oral Baclofen was not helpful. Finally, he underwent left Achilles tendon release and left tibialis anterior transfer at 7.5 years of age.

Additional clinical manifestations were feeding difficulties (slow feeding of mashed food only), failure to gain weight (FTT), hypersalivation, and delayed sleep initiation and bruxism. At his last neurologic examination (age 8.3 years), he presented with dyskinetic / hypotonic CP, GMFCS 5, stereotypic hand movements (intermittent midline hand posturing), abnormal smooth eye pursuit representing cerebral visual impairment, profound intellectual disability including deficient communication skills. Overall, the clinical phenotype is DEE, but with unusual presentation. The epilepsy is moderate and well controlled by oxcarbazepine treatment (28 mg/kg), which is uncommon for DEE. However, at the same time, development is profoundly impaired and stagnated since infancy.

### Functional analysis of Na_v_1.6^G1625R^ biophysical properties

The Na_V_1.6^G1625R^ mutation replaces a conserved glycine with arginine in the S4 segment of domain IV, introducing an extra charge between the charged arginine at position 1623 and arginine at position 1626 (Fig. 2A). To characterize the biophysical ramification of the Na_V_1.6^G1625R^ mutation, Neuro-2a cells were transiently transfected with plasmids encoding for a Na_V_1.6^WT^ or Na_V_1.6^G1625R^, together with the auxiliary subunits β1 and β2. As this cell line contains endogenous Na_V_s, we used the TTX-insensitive form of Na_V_1.6, and blocked the endogenous currents with 500 nM of TTX [5]. Whole-cell voltage-clamp recordings demonstrated lower current amplitudes of Na_V_1.6^G1625R^ compared to Na_V_1.6^WT^ (Fig. 2B, C). Examination of the effect of this mutation on its voltage dependency, demonstrated a depolarizing shift in channel activation and in the voltage dependency of channel inactivation (Fig. 2D). Measurements of the channel kinetics of inactivation showed a marked increase in the time constant of fast inactivation, leading to considerable current remaining at the end of the 20 ms voltage step (Fig. 2E, F). Consistently, Na_V_1.6^G1625R^ displayed a larger persistent current in response to a depolarizing ramp stimulus from -60 mV to 0 mV over 300 ms (Fig. 2G). Next, we examined the rate of recovery from fast inactivation, which was unaltered when the holding potential was set to -120 mV (Fig. 2H). Conversely, channel availability following repeated depolarization steps to -10 mV, from a holding potential of -70 mV, demonstrated greater availability of Na_V_1.6^G1625R^ channels. Specifically, the amplitude of Na_V_1.6^WT^ dropped by ∼40% from the first to the second depolarizing step and by ∼50% at the 9^th^. In comparison, the amplitude of Na_V_1.6^G1625R^ decreased by only 9% from the first to the second pulse and by 34% in the 9^th^ pulse (Fig. 2I, J). A similar trend was observed in the area under the curve, which reflects both the reduced peak current and the slower inactivation rate in Na_V_1.6^G1625R^ (Fig. 2K). Together, the biophysical characterization of the DEE-associated Na_V_1.6^G1625R^ mutation demonstrated changes in multiple parameters. Properties that indicated LoF included lower amplitudes and a depolarizing shift in the voltage dependency of channel activation. GoF properties included a depolarizing shift of channel inactivation, slower kinetic and fast inactivation, increased amplitude in response to gradual ramp depolarization, and augmented availability in high-frequency depolarizations.

**Fig 2.**
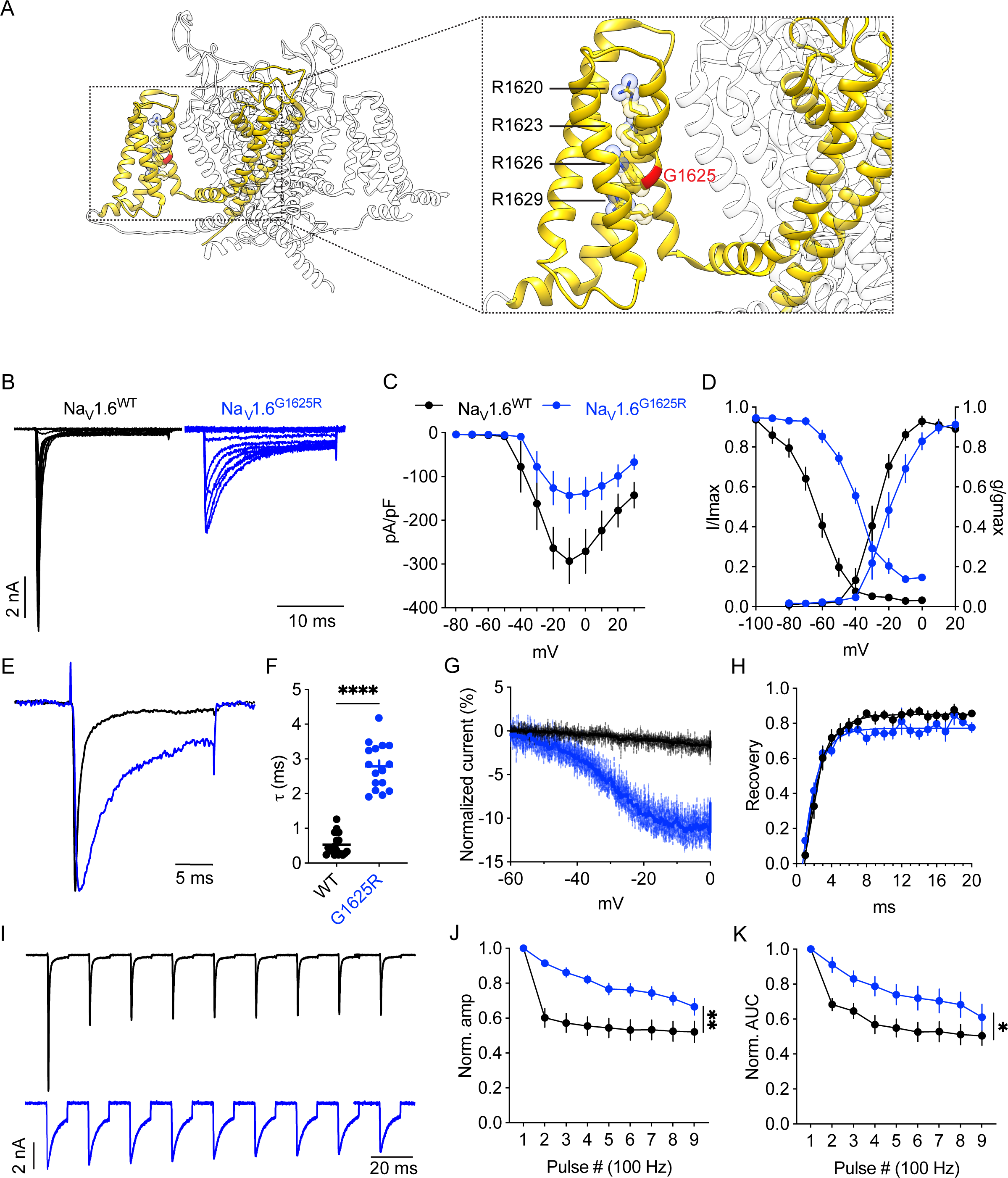
Biophysical characterization of Na_V_1.6^G1625R^ channels in Neuro-2a cells. **(A)** The location of the G1625 residue in the VSD of domain IV of Na_V_1.6 (PDB 8FHD [34]). **(B)** Representative whole-cell currents from Neuro-2a cells transfected with Na_V_1.6^WT^ or Na_V_1.6^G1625R^. **(C)** Mean current-voltage (I-V) relationships of sodium current densities**. (D)** Steady-state activation (right curves: V_1/2_: WT: -28.3 ± 2.1 mV, G1625R: -16 ± 3.6 mV, students t-test p = 0.005; Na_V_1.6^WT^ n = 18; Na_V_1.6^G1625R^ n = 15) and inactivation (left curves: V_1/2_: WT: -66.9 ± 2.3 mV, G1625R: -40 ± 1.4 mV, students t-test, p < 0.0001; Na_V_1.6^WT^ n = 17; Na_V_1.6^G1625R^ n = 10). **(E)** Normalized currents at 0 mV. **(F)** Fast inactivation time constants (τ) of WT and G1625R channels at 0mV (WT: 0.5 ± 0.007; G1625R: 3 ± 0.2, students t-test, p < 0.0001). **(G)** Averaged ramp currents (-60 to 0mV lasting 300 ms), normalized to peak currents. **(H)** Recovery from fast inactivation. **(I)** Representative traces of repetitive 10 ms depolarizations from a holding potential of -70 mV to -10 mV in 10 ms intervals. **(J)** Normalized amplitude. Mixed-model repeated measure ANOVA, p = 0.0042; Na_V_1.6^WT^ n = 11; Na_V_1.6^G1625R^ n = 8. **(K)** Normalized area under the curve. Mixed-model repeated measure ANOVA, p = 0.026.

### Reduced neuronal firing in hippocampal neurons expressing Na_V_1.6^G1625R^

Since G1625R caused both GoF and LoF changes in the biophysical properties, we further examined the functional effects of this variant on neuronal parameters and action potential firing. Current clamp recordings were performed in cultured hippocampal neurons transfected with TTX-insensitive Na_V_1.6^WT^ or Na_V_1.6^G1625R^ plasmids. The advantage of using TTX-insensitive channels in cultured neurons is that we could measure the overall firing mediated by the endogenously and exogenously expressed Na_V_s, as well as add TTX and analyze the effect of the mutation when the sodium current is mediated solely by the exogenously expressed Na_V_1.6.

In the absence of TTX in the bath solution, neurons expressing Na_V_1.6^G1625R^ did not differ from neurons expressing Na_V_1.6^WT^ in either the action potential (AP) firing or neuronal parameters (Fig. 3A-I). Conversely, in the presence of TTX, fewer action potential firing was observed for neurons expressing Na_V_1.6^G1625R^ than those expressing Na_V_1.6^WT^ (Fig. 3J). This difference was also evident in the area under the curve (AUC) for input-output relationships (Fig. 3K, L). In addition, we also observed a higher AP firing threshold in neurons expressing the variant (Fig. 3O), and a tendency for higher rheobase (Fig. 3Q). Interestingly, under these conditions, 14 out of 16 neurons expressing Na_V_1.6^WT^ exhibited AP firing, as opposed to only 5 out of 15 neurons expressing Na_V_1.6^G1625R^ (** p = 0.0032, Fisher’s exact test). To confirm the expression of exogenous Na_V_1.6, we also recorded the sodium current mediated only by transfected channels in voltage clamp mode. These recordings demonstrated reduced, yet measurable, peak inward current amplitudes in neurons expressing Na_V_1.6^G1625R^ (Fig. 3R, the minimum amplitude was 106.8 pA).

**Fig. 3:**
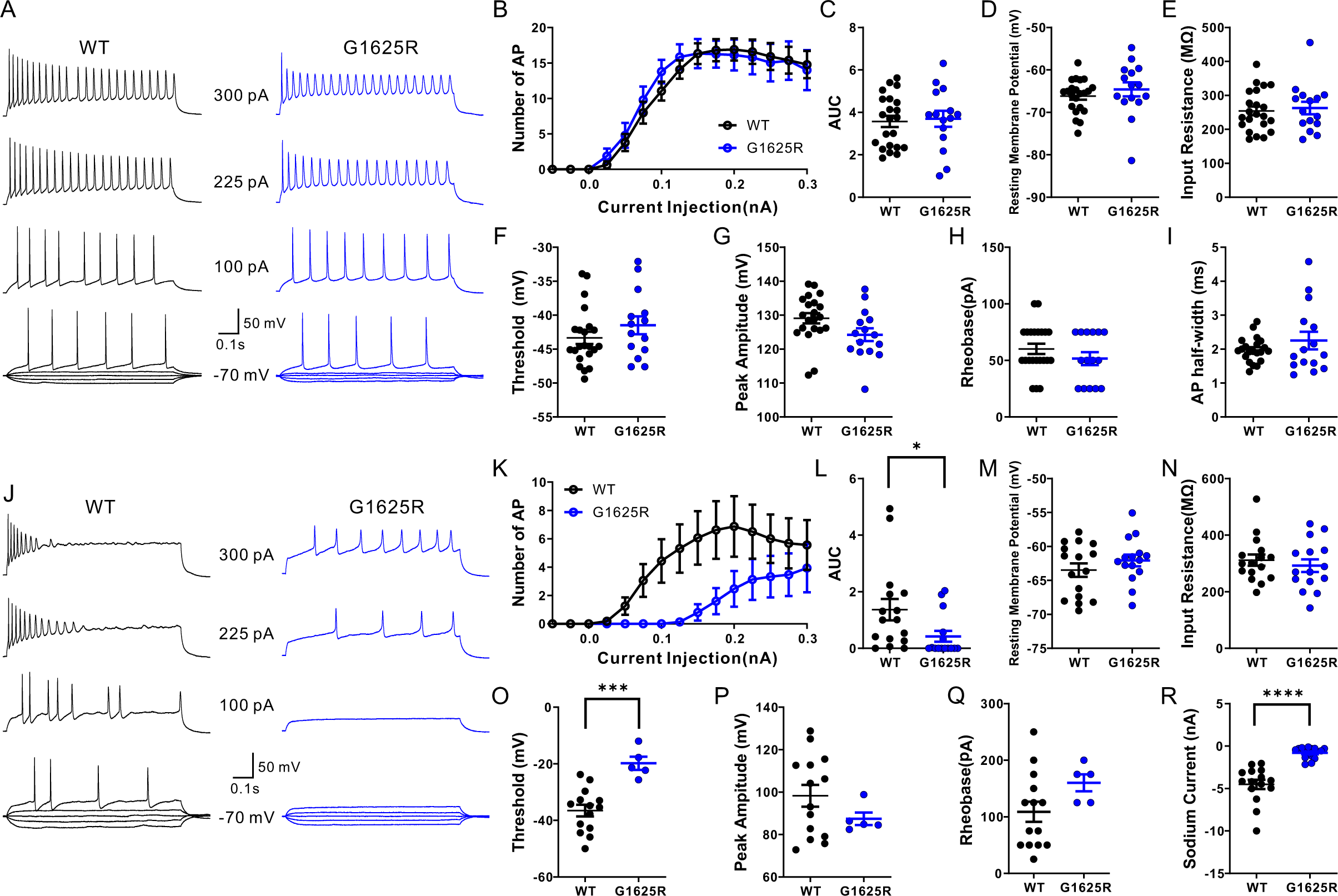
Intrinsic neuronal properties and action potential firing in neurons transfected with Na_V_1.6^WT^ or Na_V_1.6^G1625R^ in the absence or presence of TTX. **(A, J)** Representative traces of action potential firing, recorded from cultured hippocampal neurons transfected with either Na_V_1.6^WT^ or Na_V_1.6^G1625R^ in the absence (A) or presence (J) of TTX. **(B-I)** In absence of TTX, numbers of action potentials plotted versus injected current (B), area under the curve (AUC) for the input-output relationships (C), resting membrane potential (D), input resistance (E), threshold of evoked action potential (F), peak amplitude of evoked action potential (G), rheobase (H) and half-width of evoked action potential (I) were comparable between neurons with Na_V_1.6^WT^ and neurons with Na_V_1.6^G1625R^. **(K-R)** In the presence of TTX, neurons expressing Na_V_1.6^G1625R^ demonstrated decreased numbers of action potential in response to a range of injected currents (K), the area under the curve (AUC) for the input-output relationships (L), reduced inward Na^+^ current amplitudes for Na_V_1.6^G1625R^ (R), and increased threshold of evoked action potential (O). There were no differences in the resting membrane potential (M), input resistance (N), peak amplitude of evoked action potential (P), or rheobase (Q). Data are presented as mean ± SEM. Students t-test was used for normally distributed data, and the Mann-Whitney U test was used for not normally distributed data. * p<0.05; *** p<0.001;**** p<0.0001.

Overall, our results in cultured hippocampal neurons demonstrated that the intricate biophysical changes observed in our voltage clamp analyses of Na_V_1.6^G1625R^ cumulate to a LoF effect, which was mainly exemplified when the neuronal sodium currents were mediated by the transfected channels.

### Computational modeling of Na_v_1.6^G1625R^ effect on neuronal firing

Transient transfection of ion channels in cultured neurons provides a valuable framework for evaluating the functional consequences of disease-related mutations. However, this experimental paradigm has limitations, including the obligatory overexpression of WT and pathogenic variants against a background of endogenously expressed Na_V_ channels, or, in the presence of TTX, the sole reliance on the transfected channels for neuronal activity. In addition, cultured neurons show variable ion channel compositions compared to mature neurons in intact circuits *in vivo*. Thus, it may be difficult to interpret the contribution of the mutant channel to neuronal excitability without knowing the exact complementation mechanisms of endogenous channels and their possible functional interaction with the specific mutation. Conversely, computational modeling allows precise control over the channel composition, enabling systematic analysis of how specific Na_V_ channel mutations impact defined neuronal populations. Using biophysically detailed models, we can simulate mature neurons under physiologically relevant conditions, providing experimentally unattainable insights into the functional effects of Na_V_ channel mutations on neuronal excitability [35,36]. Moreover, computational modeling enables the simulation of heterozygous genetic conditions, providing a more nuanced understanding of the mutations’ effects. Here, To further explore the effects of Na_V_1.6^G1625R^ expression, we modified a well-established computational model of a layer V cortical pyramidal cell [27,28,30,37–39]. We optimized a hidden Markov model (HMM) to represent Na_V_1.6 channels using an in-house evolutionary algorithm similar to previous studies [33,40]. The HMM model parameters were fit to mimic the kinetics of the Hodgkin-Huxley (HH) formalism used originally in the model. Unlike HH models, our HMM model allows for multiple channel states, which better recapitulate the physiological channel conditions [41–43]. Notably, when simulating whole-cell voltage clamp currents, this model enabled us to recapitulate the electrophysiological recordings from Neuro-2a cells, with reduced peak current and a slower inactivation rate, as well as depolarized voltage dependency for both activation and inactivation in Na_V_1.6^G1625R^ (Fig. 4A-C). Next, we modeled the firing of neurons overexpressing Na_V_1.6^WT^ or Na_V_1.6^G1625R^, mimicking the overexpression in cultured neurons. Under these conditions, firing in response to various stimulus currents remained largely unaltered (Fig. 4D, E), similar to our data in transfected cultured hippocampal neurons (Fig. 3A-I). Next, we simulated the addition of TTX by removing the contribution of endogenous channels, which led to a marked reduction in action potential firing across all levels of current stimuli in Na_V_1.6^G1625R^ (Fig. 4F, G). The recapitulation of the cultured neuron results by the model confirmed a direct link between the pathophysiological biophysical properties and reduced neuronal excitability.

**Fig. 4.**
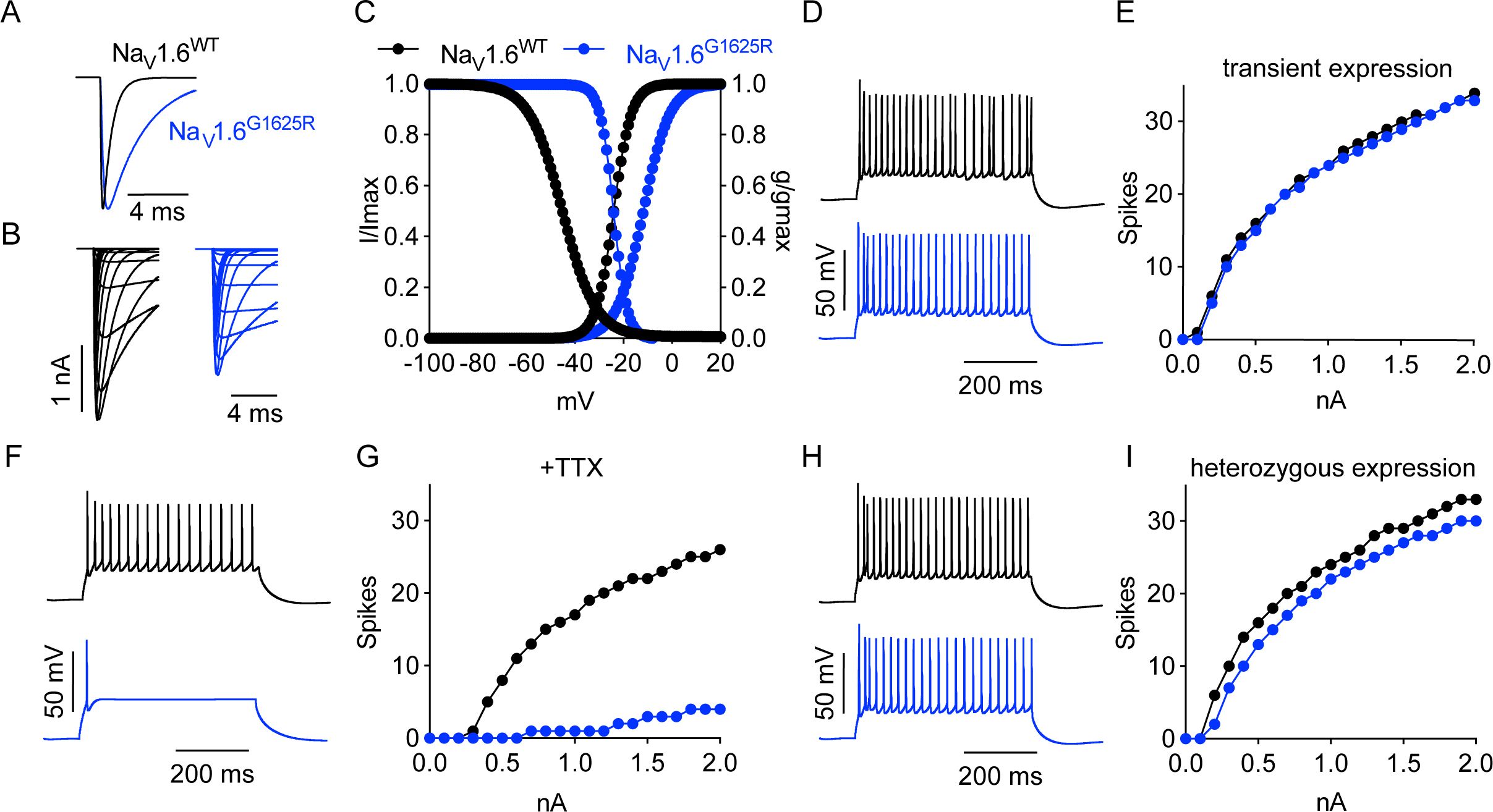
Neuronal excitability in a cortical pyramidal cell model. **(A-C)** Modeling of the biophysical properties of Na_V_1.6^WT^ (represented in black) and Na_V_1.6^G1625R^ (represented in blue). (A) Simulation of the normalized currents at 0 mV. (B) simulation of whole-cell currents. (C) the voltage dependency of activation and inactivation. **(D-I)** Simulation of neuronal firing. (D-E) Modeled firing in response to 20% overexpression of Na_V_1.6^WT^ or Na_V_1.6^G1625R^, in addition to other endogenous Na_V_ channels. Examples of modeled firing in response to injection of 1 nA current and F-I curves. (F–G) Modeled firing in response to overexpression of Na_V_1.6^WT^ or Na_V_1.6^G1625R^, without the contribution of endogenous Na_V_ channels. Examples of modeled firing in response to injection of 1 nA current and F-I curves. (H–I) Modeled firing in a patient expressing 100% Na_V_1.6^WT^ or 50% Na_V_1.6^WT^ and 50% Na_V_1.6^G1625R^. Examples of modeled firing in response to injection of 1 nA current and F-I curves.

Finally, we simulated the heterozygous condition, comprising 50% Na_V_1.6^G1625R^ and 50% Na_V_1.6^WT^, compared to the control condition with 100% Na_V_1.6^WT^ (with no overexpression of Na_V_s). This model, which mimics the genetic condition of the patient, showed a moderate reduction in the firing rate in the heterozygous state. Examination of the underlying currents during the AP, revealed that the altered properties of Na_V_1.6^G1625R^ affect the currents of other neuronal ion channels, including Na_V_1.2 currents and voltage-gated calcium channels (Supplementary Fig. 1).

Our experimental data showed a lower current amplitude in Neuro-2a and cultured neurons for Na_V_1.6^G1625R^. To test the contribution of this property to the overall LoF effect, we modeled the firing under conditions in which the current amplitudes of Na_V_1.6^WT^ and Na_V_1.6^G1625R^ are the same. Interestingly, also under these conditions, the effect was LoF, with a marked reduction in neuronal firing in the presence of TTX and a slight reduction in firing in the heterozygous state (Supplementary Fig.2).

### Functional interaction between F1588 and R1625 contributes to the shift in the voltage dependence of inactivation

To gain further molecular insights into the biophysical changes caused by the R1625 mutation, we modeled this mutant residue onto the recently solved cryo-EM structure of Na_V_1.6 [34]. This model suggested a possible atomic proximity between R1625 and F1588, allowing for the formation of a cation-π interaction between the positively charged arginine and the negatively charged electron cloud of the aromatic ring of the adjacent phenylalanine. If formed, this strong noncovalent bond may be involved in mediating some of the altered biophysical properties of Na_V_1.6^G1625R^ (Fig. 5A). To test this hypothesis, we mutated position 1588 to alanine, which has a smaller sidechain and transiently expressed these channels in Neuro-2a cells.

**Fig 5.**
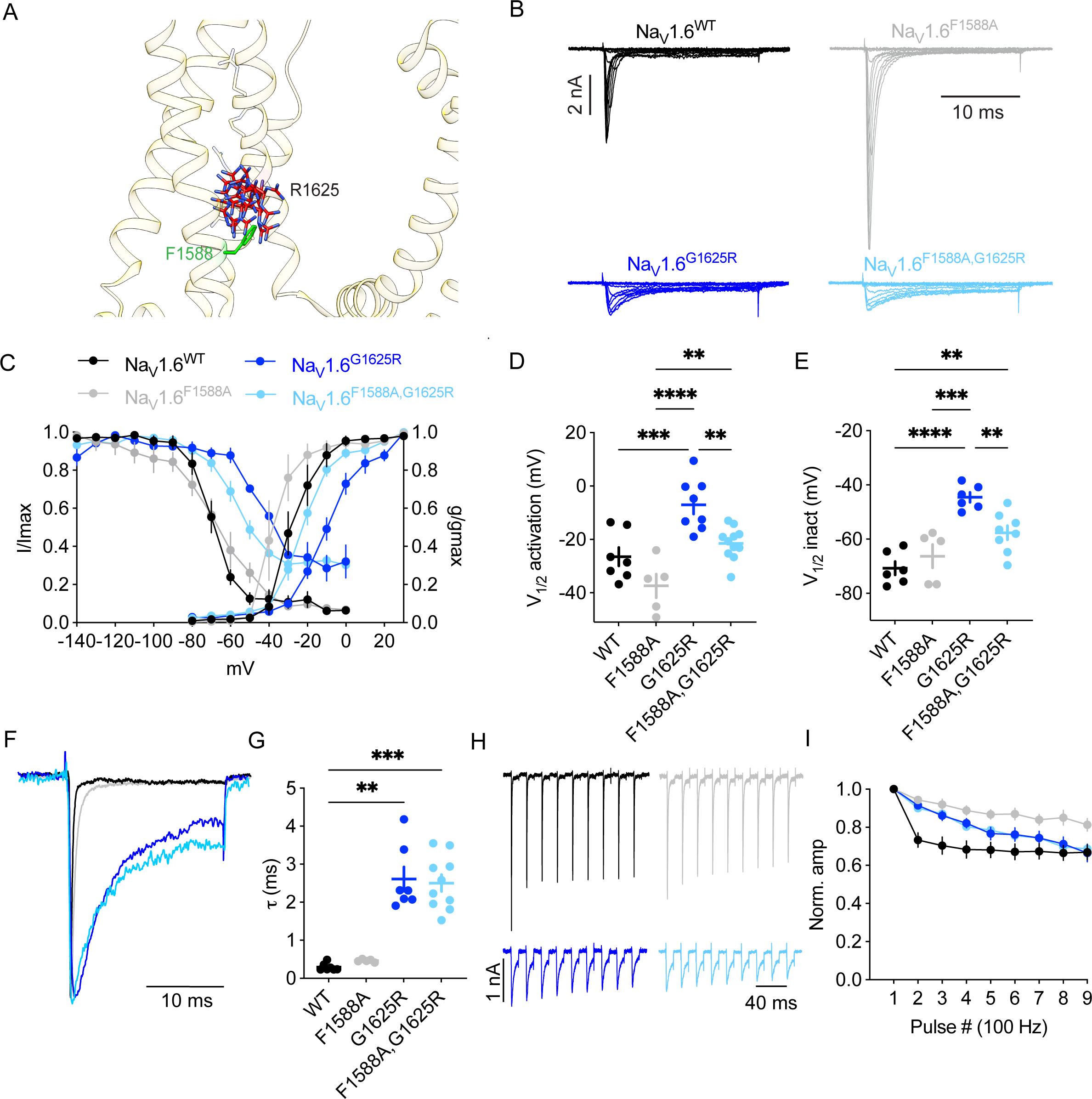
Functional interaction between R1625 and F1588. **(A)** A model of Na_V_1.6^G1625R^ structure, based on (PDB 8FHD [34]), shows the possible rotamers of R1625 and the possible interaction with F1588. **(B)** Representative current traces recorded from Neuro-2a cells, transfected with Na_V_1.6^WT^, Na_V_1.6^F1588A^, Na_V_1.6^G1625R^, and Na_V_1.6 ^F1588A,G1625R^. **(C)** Steady-state activation (right curves: V_1/2_: WT: -26.5 ± 3.4 mV, F1588A: -37.4 ± 4.4 mV; G1625R: -7 ± 3.4 mV; F1588A,G1625R: -21.43 ± 1.8 mV) and inactivation curves (left curves: V_1/2_: WT: -70.7± 2.5mV, F1588A: -66.3 ± 4.2 mV; G1625R: -44.6 ± 1.8 mV; F1588A,G1625R: - 57.6 ± 2.6 mV). **(D)** V_1/2_ of activation. Na_V_1.6^WT^ n = 7; Na_V_1.6^F1588A^ n = 5; Na_V_1.6^G1625R^ n = 8; Na_V_1.6^F1588A,G1625R^ n = 11. **(E)** V_1/2_ of inactivation. Na_V_1.6^WT^ n = 6; Na_V_1.6^F1588A^ n = 5; Na_V_1.6^G1625R^ n = 6; Na_V_1.6^F1588A/G1625R^ n = 8. One-way ANOVA, followed by Holm-Sidek multiple comparisons. **(F)** Normalized currents at 0 mV. **(G)** Inactivation time constants at 0 mV (τ). The nonparametric Kruskal-Wallis test, followed by Dunn’s multiple comparisons test. **(H)** Representative traces of repetitive 10 ms depolarizations from a holding potential of -70 mV to -10 mV in 10 ms intervals. **(J)** Normalized amplitude. ∗∗p < 0.01, ∗∗∗p < 0.001, and ∗∗∗∗p < 0.0001.

First, we analyzed the effect of the F1588A mutation. This substitution resulted in large current amplitudes that tended to activate at more hyperpolarized potentials compared to Na_V_1.6^WT^ (Fig. 5B-D: Na_V_1.6^WT^: -25.5 ± 3.4 mV, n = 7; Na_V_1.6^F1588A^: -37.4 ± 4.4 mV, n = 5; One Way ANOVA, p = 0.068). Moreover, while we did not observe changes in the voltage dependency of inactivation or the inactivation rate, Na_V_1.6^F1588A^ led to increased channel availability with repeated depolarizations at 100 Hz (Fig. 5C-I).

Next, we tested the effect of the double mutant Na_V_1.6 ^F1588A,G1625R,^, designed to dismantle the possible cation-π interaction. Compared to the single Na_V_1.6^G1625R^ mutant, the voltage dependency of activation was more hyperpolarized in the double mutant Na_V_1.6 ^F1588A,G1625R^ (Fig. 5C, D: Na_V_1.6^G1625R^: -6.99 ± 3.43 mV, n = 8; Na_V_1.6 ^F1588A,G1625R^: -21.43 ± 1.8 mV, n = 11; one way ANOVA, \*\**p* = 0.005). However, given that a minor leftward shift in the activation voltage was also observed in Na_V_1.6^F1588A^, we employed a double-mutant cycle analysis [44] to assess the degree of functional interaction between these positions. This analysis revealed a coupling free energy (ΔΔG_0_) of 0.59 ± 0.077 kcal/mol. The coupling value indicates an additive effect, suggesting a lack of functional interaction between residues R1625 and F1588 in setting the voltage dependency of channel activation.

Conversely, focusing on the voltage dependency of steady-state inactivation, theV_1/2_ values were similar between Na_V_1.6^WT^ and Na_V_1.6^F1588A^ (Fig. 5C, E: Na_V_1.6^WT^: -70.75 ± 2.5 mV, n = 6; Na_V_1.6^F1588A^: -66.27 ± 4.26 mV, n = 5; one way ANOVA, *p* = 0.3). However, when the F1588A mutation was introduced on top of Na_V_1.6^G1625R^, a significant hyperpolarizing shift was observed in the steady-state inactivation voltage (Fig. 5C, E: Na_V_1.6^G1625R^: -44.56 ± 1.84 mV, n = 6; Na_V_1.6 ^F1588A,G1625R^: -57.63 ± 2.6 mV, n = 8; one way ANOVA, \*\**p* = 0.009). Examination of the coupling free energy of voltage-dependent inactivation revealed a ΔΔG_0_ of 4.75 ± 0.088 kcal/mol, signifying a strong functional interaction between residues R1625 and F1588 in determining the voltage setting for channel inactivation.

The rate of channel inactivation was comparable between the single mutant Na_V_1.6^G1625R^ and the double mutant Na_V_1.6 ^F1588A,G1625R^. Similarly, channel availability in response to repeated depolarizations did not significantly differ between these two variants. However, it is noteworthy that the Na_V_1.6^F1588A^ mutation led to a pronounced increase in channel availability on its own, especially noticeable during the 9^th^ pulse of depolarization (Fig. 5H-I), suggesting that the effect of Na_V_1.6^G1625R^ dominated in the context of the double mutant, dampening the impact of Na_V_1.6^F1588A^.

Together, these results highlight the functional interaction between positions R1625 and F1588 and its contribution to the pathophysiological dysfunction of Na_v_1.6^G1625R^, mainly by affecting the voltage dependency of inactivation in the Na_V_1.6 mutation.

## Discussion

*SCN8A* mutations are associated with multiple neurological phenotypes, ranging from global developmental delay without epilepsy to intractable DEE. Functional analysis of missense DEE-associated mutations demonstrated a prevailing propensity for GoF mutations, associated with a hyperpolarizing shift in the voltage dependency of activation, slower inactivation kinetics, and increased persistent currents amplitude (Supplementary Table 1). These biophysical properties cumulate, in most cases, to increased neuronal firing, and result in treatment recommendations consisting of sodium channel blockers [3,45]. However, some patients with *SCN8A*-DEE were found to harbor frameshift stop-gain or canonical splice-site mutations, leading to LoF [3]. Thus, while *SCN8A*-DEE-causing mutations often cause gain of channel function, some mutations may have a more complex functional outcome. Similar to other canonical GoF DEE-causing mutations, Na_V_1.6^G1625R^ caused a depolarizing shift in the voltage dependency of inactivation, slower inactivation kinetics, augmented currents in response to a slow ramp-like stimulus, and enhanced availability with repeated depolarizations (Fig. 2). However, we also observed several LoF characteristics, presented as a lower amplitude in Neuro-2a cells (Fig. 2) and cultured hippocampal neurons (Fig. 3), and a depolarizing shift in the voltage dependency of activation (Fig, 2, 5). When transfected to cultured neurons, in the presence of TTX, the intricate functional alterations of Na_V_1.6^G1625R^ led to reduced neuronal firing (Fig. 3). However, when modeled to simulate heterozygosity, mimicking the patients’ genetic expression profile, we observed a mild reduction in firing, indicating an overall moderate LoF (Fig. 4).

What might be the cause for this overall mild LoF effect? One possibility is the reduced amplitude. Indeed, reduced current amplitude and firing were observed in other Na_V_1.6 DEE-associated mutations, V216D [14] and R223G [5]. However, testing this possibility using our computational model deemed the reduced amplitude as a less likely cause, as reduced firing was observed even when the amplitudes of Na_V_1.6^WT^ and Na_V_1.6^G1625R^ were set to similar values (Supplementary Fig. 2). Another possibility is the depolarized shift in the voltage dependency of activation (Fig. 2, Supplementary Table 1). Although we could not directly test that in the model, as changing the activation voltage also affected the inactivation voltage dependency and the persistent currents (data not shown), the N1768D mutation, which also features depolarized voltage dependency of activation, was previously shown to display an increased firing [17,18]. Thus, it is possible that the reduced firing of neurons expressing the G1625R mutation, especially in the presence of TTX, is driven by the convergence of multiple unique biophysical changes.

Mutations in homologous positions in other voltage-gated sodium channels were reported, supporting the functional importance of this position. One of these, the de novo *SCN5A* G1631D mutation, was identified in two patients with neonate severe arrhythmia and long QT syndrome. Functional analysis of this mutation demonstrated GoF properties that were effectively treated with sodium channel blockers [46]. While the biophysical characterization demonstrated properties reminiscent of Na_V_1.6^G1625R^, the *SCN5A* G1613D mutation was associated with increased open probability and a slower recovery from inactivation, which pushed the overall properties to a GoF [46]. Mutations in other Na_V_ channel genes were also reported, but to the best of our knowledge, with no functional testing. Interestingly, clinical prediction tools were inconclusive regarding the functional ramifications of mutations in this glycine residue (Supplementary Table 2), supporting the importance of functional analysis.

Other pathogenic *SCN8A* mutations also showed compound biophysical changes. For instance, the DEE-associated *SCN8A* R223G mutation was shown to cause GoF properties, including a hyperpolarizing shift in the voltage dependency of activation [15] and increased ramp currents [6]. Concurrently, LoF features for this mutation involved a reduced current amplitude [6,15] and a hyperpolarizing shift in the voltage dependency of inactivation [15]. The overall effect was increased firing in the absence of TTX, yet a marked reduction in the number of action potentials (although with reduced rheobase) in the presence of TTX when the firing is forced solely by the exogenously expressed Na_V_1.6 [15]. Likewise, the *SCN8A-* DEE associated N1768D mutation demonstrated mixed effects in expression systems with reduced current amplitude, and depolarized shift in the voltage dependency of channel activation, but with increased persistent current and increased neuronal firing when transfected to cultured hippocampal neurons [17]. Nevertheless, recordings from *Scn8a*^WT/N1768D^ mice revealed compound effects with GoF or LoF characteristics [18,47–49]. Along this line, studies of mouse models carrying *Scn8a* GoF mutations further supported the complexity of the neuronal mechanism in DEE-associated *SCN8A* mutations, demonstrating diverse effects on neuronal activity in different brain regions, neuronal subtypes, and developmental stages [18,47–51].

The G1625R mutation is located in the voltage sensor of domain IV, adding another charge between R2 and R3 in the fourth transmembrane segment (S4) of domain IV. This region was shown before to contribute to the coupling between activation and inactivation and to affect the inactivation time constant [52–55]. The G1625R substitution is uncommon in the realm of disease-causing sodium channelopathies as it adds another charge to the already charged S4, while most of the homologous mutations at this position in other Na_V_ channels result in the neutralization of one of the charged S4 arginine [56,57]. Similarly, studies that examine the functional effect of S4 domain IV also employed charge neutralization [52,53,55], while the functional effect of charge additions was less explored. Nevertheless, similar to the data presented here, charge perturbation along S4 domain IV dramatically modulated the kinetic of Na_V_1.6 channel inactivation.

Modeling the R1625 mutation on the structure of Na_V_1.6 indicated that the charged arginine side chain may be in close proximity to F1588, located in S3 domain IV, enabling the formation of a cation-π interaction. Cation-π interactions were shown to mediate the binding of local anesthetics and toxins to voltage-gated sodium channels [58], and contribute to the functional effect of disease-related mutations in HCN channels and TRPV [59,60]. Our double mutant cycle analysis suggested that disruption of this cation-π interaction, via the F1588A substitution, corrected the depolarized shift in the voltage dependency of inactivation caused by the G1625R mutation, indicating the functional interaction between these positions (Fig. 5). Yet, no function interaction was observed between R1625 and F1588 in affecting the channel availability, reduced current amplitude, or the slower rate of inactivation (Fig. 5). Interestingly, The F1588A mutation showed GoF properties by itself, with a marked increase in channel availability with repeated depolarization, a tendency for hyperpolarization shift in the voltage dependency of activation, and a tendency for a slower rate of inactivation (Fig. 5). This is in agreement with functional studies of the homologous Na_V_1.2 mutation, F1597L, which was identified in a patient with epilepsy of infancy with migrating focal seizures. Indeed, biophysical characterization of Na_V_1.2^F1597L^ demonstrated a hyperpolarizing shift of the activation curve, slow inactivation rate, and fast recovery from fast inactivation [61]. In Na_V_1.6, the F1588L mutation was identified in 2 patients with intermediate epilepsy [62], but their functional analysis was not performed.

Together, our analysis demonstrated that the Na_v_1.6^G1625R^ leads to convoluted GoF and LoF biophysical properties that result in a mild reduction in neuronal firing and are partially related to aberrant functional interaction within the voltage sensor domain of domain IV.

## Acknowledgments

We acknowledge the financial support of The Israel Science Foundation (1454/17, 214/22 to MR, and 1653/21 to YH), The German Research Foundation (DFG Research Unit FOR-2715, grant Le1030/15-2), The Federal Ministry of Education and Research (BMBF, Treat-ION, 01GM2210A), and The Hartwell Foundation through an Individual Biomedical Research (RBS).

Additional support was from The Claire and Amedee Maratier Institute for the Study of Blindness and Visual Disorders, Faculty of Medicine, Tel-Aviv University (MR, MG), and The Stolz Foundation Faculty of Medicine, Tel-Aviv University (MR), the Kahn Foundation’s Orion project, Tel Aviv Sourasky Medical Center, Israel (MG.). The Israel Cancer Research Fund grant 19202 (MG), the Israel Cancer Association grants 20230029 (YH and MG).

This work was performed in partial fulfillment of the requirements for a Ph.D. degree of SQ. We also wish to thank Dr. Mahmoud Koko for his kind help with DNA plasmids and mutagenesis, and Prof. Maya Schuldiner for introducing us to this project and initiating this fruitful collaboration.

## Authors contribution

Conception and design of the study: SQ, NZ, TAF, YH, HL, YL, RBS, MR. Acquisition and analysis of data: SQ, NZ, TAF, MB, PM, YP, MG, YH, HB. Drafting a significant portion of the manuscript or figures: SQ, NZ, TAF, HB, YL, RBS, MR. All authors have read and approved the final version of the manuscript.

## Declaration of interests

The authors declare that the research was conducted in the absence of any commercial or financial relationships that could be construed as a potential conflict of interest.

## Notes

### Competing Interest Statement

The authors have declared no competing interest.

### Summary of Updates

we revised the title, reformated the abstract, and corrected several typos in the figures

## References

[1] Mantegazza M, Cestèle S, Catterall WA. Sodium channelopathies of skeletal muscle and brain. Physiol Rev 2021;101:1633–89. 10.1152/physrev.00025.2020.

[2] Talwar D, Hammer MF. *SCN8A* epilepsy, developmental encephalopathy, and related disorders. Pediatr Neurol 2021;122:76–83. 10.1016/J.PEDIATRNEUROL.2021.06.011.

[3] Johannesen KM, Liu Y, Koko M, Gjerulfsen CE, Sonnenberg L, Schubert J, et al. Genotype-phenotype correlations in *SCN8A*-related disorders reveal prognostic and therapeutic implications. Brain 2022;145:2991–3009. 10.1093/brain/awab321.

[4] Hack JB, Horning K, Short DMJ, Schreiber JM, Watkins JC, Hammer MF. Distinguishing loss-of-function and gain-of-function *SCN8A* variants using a random forest classification model trained on clinical features. Neurol Genet 2023;9:e200060. 10.1212/NXG.0000000000200060.

[5] Liu Y, Schubert J, Sonnenberg L, Helbig KL, Hoei-Hansen CE, Koko M, et al. Neuronal mechanisms of mutations in *SCN8A* causing epilepsy or intellectual disability. Brain 2019;142:376–90. 10.1093/brain/awy326.

[6] de Kovel CGF, Meisler MH, Brilstra EH, van Berkestijn FMC, Slot R van t., van Lieshout S, et al. Characterization of a de novo *SCN8A* mutation in a patient with epileptic encephalopathy. Epilepsy Res 2014;108:1511–8. 10.1016/J.EPLEPSYRES.2014.08.020.

[7] Estacion M, O’Brien JE, Conravey A, Hammer MF, Waxman SG, Dib-Hajj SD, et al. A novel de novo mutation of *SCN8A* (Na_V_1.6) with enhanced channel activation in a child with epileptic encephalopathy. Neurobiol Dis 2014;69:117–23. 10.1016/J.NBD.2014.05.017.

[8] Pan Y, Cummins TR, Forsythe I, Marra V. Distinct functional alterations in *SCN8A* epilepsy mutant channels. J Physiol 2020;598:381–401. 10.1113/JP278952.

[9] Blanchard MG, Willemsen MH, Walker JB, Dib-Hajj SD, Waxman SG, Jongmans MCJ, et al. De novo gain-of-function and loss-of-function mutations of *SCN8A* in patients with intellectual disabilities and epilepsy. J Med Genet 2015;52:330–7. 10.1136/JMEDGENET-2014-102813.

[10] Barker BS, Ottolini M, Wagnon JL, Hollander RM, Meisler MH, Patel MK. The *SCN8A* encephalopathy mutation p.Ile1327Val displays elevated sensitivity to the anticonvulsant phenytoin. Epilepsia 2016;57:1458–66. 10.1111/EPI.13461.

[11] Guo Q bei, Zhan L, Xu H yan, Gao Z bing, Zheng Y ming. *SCN8A* epileptic encephalopathy mutations display a gain-of-function phenotype and divergent sensitivity to antiepileptic drugs. Acta Pharmacol Sin 2022 4312 2022;43:3139–48. 10.1038/s41401-022-00955-x.

[12] Zaman T, Abou Tayoun A, Goldberg EM. A single[center *SCN8A*[related epilepsy cohort: clinical, genetic, and physiologic characterization. Ann Clin Transl Neurol 2019;6:1445. 10.1002/ACN3.50839.

[13] Wagnon JL, Barker BS, Hounshell JA, Haaxma CA, Shealy A, Moss T, et al. Pathogenic mechanism of recurrent mutations of *SCN8A* in epileptic encephalopathy. Ann Clin Transl Neurol 2016;3:114. 10.1002/ACN3.276.

[14] Solé L, Wagnon JL, Tamkun MM. Functional analysis of three Na_V_1.6 mutations causing early infantile epileptic encephalopathy. Biochim Biophys Acta - Mol Basis Dis 2020;1866:165959. 10.1016/J.BBADIS.2020.165959.

[15] Liu Y, Koko M, Lerche H. A *SCN8A* variant associated with severe early onset epilepsy and developmental delay: Loss- or gain-of-function? Epilepsy Res 2021;178:106824. 10.1016/J.EPLEPSYRES.2021.106824.

[16] Poulin H, Chahine M, Toth K, Young S, Poulin H, Chahine M, et al. R1617Q epilepsy mutation slows Na_V_1.6 sodium channel inactivation and increases the persistent current and neuronal firing. J Physiol 2021;599:1651–64. 10.1113/JP280838.

[17] Veeramah KR, O’Brien JE, Meisler MH, Cheng X, Dib-Hajj SD, Waxman SG, et al. De novo pathogenic *SCN8A* mutation identified by whole-genome sequencing of a family quartet affected by infantile epileptic encephalopathy and SUDEP. Am J Hum Genet 2012;90:502–10. 10.1016/J.AJHG.2012.01.006.

[18] Wengert ER, Saga AU, Panchal PS, Barker BS, Patel MK. Prax330 reduces persistent and resurgent sodium channel currents and neuronal hyperexcitability of subiculum neurons in a mouse model of *SCN8A* epileptic encephalopathy. Neuropharmacology 2019;158:107699. 10.1016/j.neuropharm.2019.107699.

[19] Tidball AM, Lopez-Santiago LF, Yuan Y, Glenn TW, Margolis JL, Clayton Walker J, et al. Variant-specific changes in persistent or resurgent sodium current in *SCN8A*-related epilepsy patient-derived neurons. Brain 2020;143:3025–40. 10.1093/brain/awaa247.

[20] Encinas AC, Watkins JC, Longoria IA, Johnson JP, Hammer MF. Variable patterns of mutation density among Na_V_1.1, Na_V_1.2 and Na_V_1.6 point to channel-specific functional differences associated with childhood epilepsy. PLoS One 2020;15. 10.1371/JOURNAL.PONE.0238121.

[21] Firth H V., Richards SM, Bevan AP, Clayton S, Corpas M, Rajan D, et al. DECIPHER: Database of chromosomal imbalance and phenotype in humans using ensembl resources. Am J Hum Genet 2009;84:524–33. 10.1016/J.AJHG.2009.03.010.

[22] Boßelmann CM, Hedrich UBS, Lerche H, Pfeifer N. Predicting functional effects of ion channel variants using new phenotypic machine learning methods. PLOS Comput Biol 2023;19:e1010959. 10.1371/JOURNAL.PCBI.1010959.

[23] Heyne HO, Baez-Nieto D, Iqbal S, Palmer DS, Brunklaus A, May P, et al. Predicting functional effects of missense variants in voltage-gated sodium and calcium channels. Sci Transl Med 2020;12:6848. 10.1126/SCITRANSLMED.AAY6848/SUPPL_FILE/AAY6848_TABLE_S6.TXT.

[24] Brunklaus A, Feng T, Brünger T, Perez-Palma E, Heyne H, Matthews E, et al. Gene variant effects across sodium channelopathies predict function and guide precision therapy 2022;145:4275–86. 10.1093/BRAIN/AWAC006.

[25] Nissenkorn A, Almog Y, Adler I, Safrin M, Brusel M, Marom M, et al. In vivo, in vitro and in silico correlations of four de novo *SCN1A* missense mutations. PLoS One 2019;14:e0211901. 10.1371/journal.pone.0211901.

[26] Meisel E, Dvir M, Haitin Y, Giladi M, Peretz A, Attali B. KCNQ1 channels do not undergo concerted but sequential gating transitions in both the absence and the presence of KCNE1 protein. J Biol Chem 2012;287:34212–24. 10.1074/JBC.M112.364901.

[27] Spratt PWE, Alexander RPD, Ben-Shalom R, Sahagun A, Kyoung H, Keeshen CM, et al. Paradoxical hyperexcitability from Na_V_1.2 sodium channel loss in neocortical pyramidal cells. Cell Rep 2021;36:109483. 10.1016/j.celrep.2021.109483.

[28] Hallermann S, De Kock CP, Stuart GJ, Kole MH. State and location dependence of action potential metabolic cost in cortical pyramidal neurons. Nat Neurosci 2012;15:1007–14. 10.1038/nn.3132.

[29] Ramaswamy S, Markram H. Anatomy and physiology of the thick-tufted layer 5 pyramidal neuron. Front Cell Neurosci 2015;9:144541. 10.3389/FNCEL.2015.00233/BIBTEX.

[30] Ben-Shalom R, Keeshen CM, Berrios KN, An JY, Sanders SJ, Bender KJ. Opposing effects on Na_V_1.2 function underlie differences between *SCN2A* variants observed in Individuals with autism spectrum disorder or infantile seizures. Biol Psychiatry 2017;82:224–32. 10.1016/J.BIOPSYCH.2017.01.009.

[31] Schmidt-Hieber C, Bischofberger J. Fast sodium channel gating supports localized and efficient axonal action potential initiation. J Neurosci 2010;30:10233–42. 10.1523/JNEUROSCI.6335-09.2010.

[32] Van Geit W, Gevaert M, Chindemi G, Rössert C, Courcol JD, Muller EB, et al. BluePyOpt: Leveraging open source software and cloud infrastructure to optimise model parameters in neuroscience. Front Neuroinform 2016;10:195490. 10.3389/FNINF.2016.00017/BIBTEX.

[33] Ladd A, Kim KG, Balewski J, Bouchard K, Ben-Shalom R. Scaling and benchmarking an evolutionary algorithm for constructing biophysical neuronal models. Front Neuroinform 2022;16:882552. 10.3389/fninf.2022.882552.

[34] Fan X, Huang J, Jin X, Yan N. Cryo-EM structure of human voltage-gated sodium channel Na_V_1.6. Proc Natl Acad Sci U S A 2023;120:e2220578120. 10.1073/pnas.2220578120.

[35] Nandi A, Chartrand T, Van Geit W, Buchin A, Yao Z, Lee SY, et al. Single-neuron models linking electrophysiology, morphology, and transcriptomics across cortical cell types. Cell Rep 2022;40:111176. 10.1016/j.celrep.2022.111176.

[36] Haufler D, Ito S, Koch C, Arkhipov A. Simulations of cortical networks using spatially extended conductance-based neuronal models. J Physiol 2023;601:3123–39. 10.1113/JP284030.

[37] Echevarria-Cooper DM, Hawkins NA, Misra SN, Huffman AM, Thaxton T, Thompson CH, et al. Cellular and behavioral effects of altered Na_V_1.2 sodium channel ion permeability in *Scn2a* K1422E mice. Hum Mol Genet 2022;31:2964–88. 10.1093/hmg/ddac087.

[38] Tamura S, Nelson AD, Spratt PWE, Kyoung H, Zhou X, Li Z, et al. CRISPR activation rescues abnormalities in *SCN2A* haploinsufficiency-associated autism spectrum disorder. BioRxiv 2022:2022.03.30.486483. 10.1101/2022.03.30.486483.

[39] Spratt PWE, Ben-Shalom R, Keeshen CM, Burke KJ, Clarkson RL, Sanders SJ, et al. The autism-associated gene *Scn2a* contributes to dendritic excitability and synaptic function in the prefrontal cortex. Neuron 2019;103:673–685.e5. 10.1016/J.NEURON.2019.05.037.

[40] Ben-Shalom R, Aviv A, Razon B, Korngreen A. Optimizing ion channel models using a parallel genetic algorithm on graphical processors. J Neurosci Methods 2012;206:183–94. 10.1016/J.JNEUMETH.2012.02.024.

[41] Clerx M, Beattie KA, Gavaghan DJ, Mirams GR. Four ways to fit an ion channel model. Biophys J 2019;117:2420–37. 10.1016/J.BPJ.2019.08.001.

[42] Lampert A, Korngreen A. Markov modeling of ion channels: implications for understanding disease. Prog Mol Biol Transl Sci 2014;123:1–21. 10.1016/B978-0-12-397897-4.00009-7.

[43] Milescu LS, Yamanishi T, Ptak K, Mogri MZ, Smith JC. Real-time kinetic modeling of voltage-gated ion channels using dynamic clamp. Biophys J 2008;95:66–87. 10.1529/biophysj.107.118190.

[44] Horovitz A. Double-mutant cycles: a powerful tool for analyzing protein structure and function. Fold Des 1996;1:R121–6. 10.1016/S1359-0278(96)00056-9.

[45] Peng BW, Tian Y, Chen L, Duan LF, Wang XY, Zhu HX, et al. Genotype-phenotype correlations in *SCN8A*-related epilepsy: a cohort study of Chinese children in southern China. Brain 2022;145:e24–7. 10.1093/BRAIN/AWAC038.

[46] Wang DW, Crotti L, Shimizu W, Pedrazzini M, Cantu F, De Filippo P, et al. Malignant perinatal variant of long-QT syndrome caused by a profoundly dysfunctional cardiac sodium channel. Circ Arrhythm Electrophysiol 2008;1:370. 10.1161/CIRCEP.108.788349.

[47] Lopez-Santiago LF, Yuan Y, Wagnon JL, Hull JM, Frasier CR, O’Malley HA, et al. Neuronal hyperexcitability in a mouse model of *SCN8A* epileptic encephalopathy. Proc Natl Acad Sci U S A 2017;114:2383–8. 10.1073/pnas.1616821114.

[48] Baker EM, Thompson CH, Hawkins NA, Wagnon JL, Wengert ER, Patel MK, et al. The novel sodium channel modulator GS-458967 (GS967) is an effective treatment in a mouse model of *SCN8A* encephalopathy. Epilepsia 2018;59:1166–76. 10.1111/EPI.14196.

[49] Wengert ER, Miralles RM, Wedgwood KCA, Wagley PK, Strohm SM, Panchal PS, et al. Somatostatin-positive interneurons contribute to seizures in *SCN8A* epileptic encephalopathy. J Neurosci 2021;41:9257–73. 10.1523/JNEUROSCI.0718-21.2021.

[50] Bunton-Stasyshyn RKA, Wagnon JL, Wengert ER, Barker BS, Faulkner A, Wagley PK, et al. Prominent role of forebrain excitatory neurons in *SCN8A* encephalopathy. Brain 2019;142:362. 10.1093/BRAIN/AWY324.

[51] Ottolini M, Barker BS, Gaykema RP, Meisler MH, Patel MK. Aberrant sodium channel currents and hyperexcitability of medial entorhinal cortex neurons in a mouse model of *SCN8A* encephalopathy. J Neurosci 2017;37:7643–55. 10.1523/JNEUROSCI.2709-16.2017.

[52] Chen LQ, Santarelli V, Horn R, Kallen RG. A unique role for the S4 segment of domain 4 in the inactivation of sodium channels. J Gen Physiol 1996;108:549. 10.1085/JGP.108.6.549.

[53] Mitrovic N, George AL, Horn R, Horn D. Role of domain 4 in sodium channel slow inactivation. J Gen Physiol 2000;115:707–18. 10.1085/JGP.115.6.707.

[54] Osteen JD, Sampson K, Iyer V, Julius D, Bosmans F. Pharmacology of the Na_V_1.1 domain IV voltage sensor reveals coupling between inactivation gating processes. Proc Natl Acad Sci U S A 2017;114:6836–41. 10.1073/PNAS.1621263114/SUPPL_FILE/PNAS.201621263SI.PDF.

[55] Capes DL, Goldschen-Ohm MP, Arcisio-Miranda M, Bezanilla F, Chanda B. Domain IV voltage-sensor movement is both sufficient and rate limiting for fast inactivation in sodium channels. J Gen Physiol 2013;142:101. 10.1085/JGP.201310998.

[56] Nakajima T, Kaneko Y, Dharmawan T, Kurabayashi M. Role of the voltage sensor module in Na_V_ domain IV on fast inactivation in sodium channelopathies: The implication of closed-state inactivation. Channels 2019;13:331–43. 10.1080/19336950.2019.1649521.

[57] Menezes LFS, Sabiá Júnior EF, Tibery DV, Carneiro L dos A, Schwartz EF. Epilepsy-related voltage-gated sodium channelopathies: A review. Front Pharmacol 2020;11:554390. 10.3389/FPHAR.2020.01276/BIBTEX.

[58] Infield DT, Rasouli A, Galles GD, Chipot C, Tajkhorshid E, Ahern CA. Cation-π interactions and their functional roles in membrane proteins. J Mol Biol 2021;433:167035. 10.1016/J.JMB.2021.167035.

[59] Teng J, Anishkin A, Kung C, Blount P. Human mutations highlight an intersubunit cation–π bond that stabilizes the closed but not open or inactivated states of TRPV channels. Proc Natl Acad Sci U S A 2019;116:9410–6. 10.1073/PNAS.1820673116/SUPPL_FILE/PNAS.1820673116.SM03.AVI.

[60] Hung A, Forster IC, Mckenzie CE, Berecki G, Petrou S, Kathirvel A, et al. Biophysical analysis of an HCN1 epilepsy variant suggests a critical role for S5 helix Met-305 in voltage sensor to pore domain coupling. Prog Biophys Mol Biol 2021;166:156–72. 10.1016/J.PBIOMOLBIO.2021.07.005.

[61] Wolff M, Johannesen KM, Hedrich UBS, Masnada S, Rubboli G, Gardella E, et al. Genetic and phenotypic heterogeneity suggest therapeutic implications in *SCN2A*-related disorders. Brain 2017;140:1316–36. 10.1093/BRAIN/AWX054.

[62] Johannesen KM, Gardella E, Encinas AC, Lehesjoki AE, Linnankivi T, Petersen MB, et al. The spectrum of intermediate *SCN8A*-related epilepsy. Epilepsia 2019;60:830–44. 10.1111/EPI.14705.

