## Supplementary material for "Complex biophysical changes and reduced neuronal firing in an *SCN8A* variant associated with developmental delay and epilepsy"

| Mutation | Amplitude | $V_{1/2}$ activation | $V_{1/2}$ inactivation | Tau inactivation | Persistent / ramp currents | Firing in cultured neurons |
|---|---|---|---|---|---|---|
| G214D (Solé et al., 2020) | No change | Hyperpolarized | No change | | | Increased spontaneous firing at -50 mV, hyperpolarized threshold, reduced firing in response to ramp stimulation |
| N215D (Solé et al., 2020) | No change | Hyperpolarized | No change | | | Increased spontaneous firing, hyperpolarized threshold, reduced firing in response to ramp stimulation |
| V216D (Solé et al., 2020) | Reduced | Hyperpolarized | Hyperpolarized | | | Increased spontaneous firing at -80 mV, but reduced at -50 mV, hyperpolarized threshold, reduced firing in response to ramp stimulation |
| R223G (de Kovel et al., 2014; Liu et al., 2021) | Reduced | Hyperpolarized | No change | No change | No change / increased | Increased firing without TTX but reduced firing as well as reduced rheobase with TTX |
| T767I (Estacion et al., 2014; Pan et al., 2020) | Reduced | Hyperpolarized | No change | No change | Increased | Increased (model) |
| L840P (Johannesen et al., 2022) | No change | Hyperpolarized | No change | No change | No change | Increased (with TTX) |
| F846S (Johannesen et al., 2022) | No change | Hyperpolarized | No change | No change | No change | Increased (with and w/o TTX) |
| R850Q (Pan et al., 2020) | No change | Hyperpolarized | No change | No change | Increased | Increased (model) |
| N984K (Blanchard et al., 2015) | No change | Hyperpolarized | No change | No change | No change | |
| I1327V (Barker et al., 2016) | No change | Hyperpolarized | Depolarized | Slower | | |
| N1466T (Guo et al., 2022) | No change | No change | No change | No change | Increased | |
| N1466K (Guo et al., 2022) | No change | No change | No change | Slower | Increased | |
| G1475R (Liu et al., 2019; Zaman et al., 2019) | No change | No change | No change / depolarization | Slower | Increased | Increased (with TTX) |
| A1491V (Zaman et al., 2019) | No change | Hyperpolarized | Depolarized | Slower | Increased | |
| R1617Q (Wagnon et al., 2016; Pan et al., 2020; Poulin et al., 2021) | No change | No change / hyperpolarization | No change / depolarization | Slower | Increased | Increased (model and w/o TTX) |
| **G1625R** | **Reduced** | **Depolarized** | **Depolarized** | **Slower** | **Increased** | **Reduced (with TTX)** |
| M1760I (Liu et al., 2019) | No change | Hyperpolarized | No change | Slower | Increased | Increased (with and w/o TTX) |
| N1768D (Veeramah et al., 2012; Wengert et al., 2019) | No change / Reduced | No change / Depolarized | No change / Depolarized | | Increased | Increased (with and w/o TTX) |
| R1872W (Wagnon et al., 2016; Liu et al., 2019; Wengert et al., 2021) | No change / increased | Hyperpolarized | No change | Slower | Increased | Increased (with and w/o TTX) |
| R1872Q (Wagnon et al., 2016; Pan et al., 2020) | Increased | Hyperpolarized | Depolarized | Slower | Increased | Increased (model) |
| R1872L (Wagnon et al., 2016; Zaman et al., 2019; Tidball et al., 2020) | No change | Hyperpolarized | No change | Slower | Increased | Increased bursting |
| N1877S (Johannesen et al., 2022) | No change | No change | Depolarized | Slower | No change | Increased (with and w/o TTX) |

Supplementary Table 1. Comparison of the biophysical properties of functionally characterized *SCN8A* variants that can cause DEE.

| Gene | Variant | Clinical description | Functional analysis | SCN variant prediction tools | |
|---|---|---|---|---|---|
| | | | | (Boßelmann et al., 2023) | (Heyne et al., 2020) |
| *SCN1A* | G1644D | Epilepsy and developmental delay (Landrum et al., 2018; Lindy et al., 2018) | | 84.4% LoF | 76% GoF |
| *SCN2A* | G1634V | Ohtahara syndrome(Sanders et al., 2018) | | 61.4% LoF | 67% GoF |
| *SCN2A* | G1634D | episodic ataxia (Reynolds et al., 2020) | | 57.5% LoF | 76% GoF |
| *SCN4A* | G1456E | Paramyotonia congenita of von Eulenburg (Sasaki et al., 1999) | | 73.4% GoF | 74% GoF |
| *SCN4A* | G1456W | Hypokalemic periodic paralysis type 2 (Landrum et al., 2018) | | 58.4% LoF | 67% GoF |
| *SCN5A* | G1631D | Neonate severe arrhythmia and long QT syndrome (Wang et al., 2008) | GoF (Wang et al., 2008) | GoF, part of the training data | 76% GoF |
| *SCN8A* | G1625R | DEE | Figs. 2-4 | 67% GoF | 67% GoF |

Supplementary Table 2. Homologous mutations in other sodium channel genes.

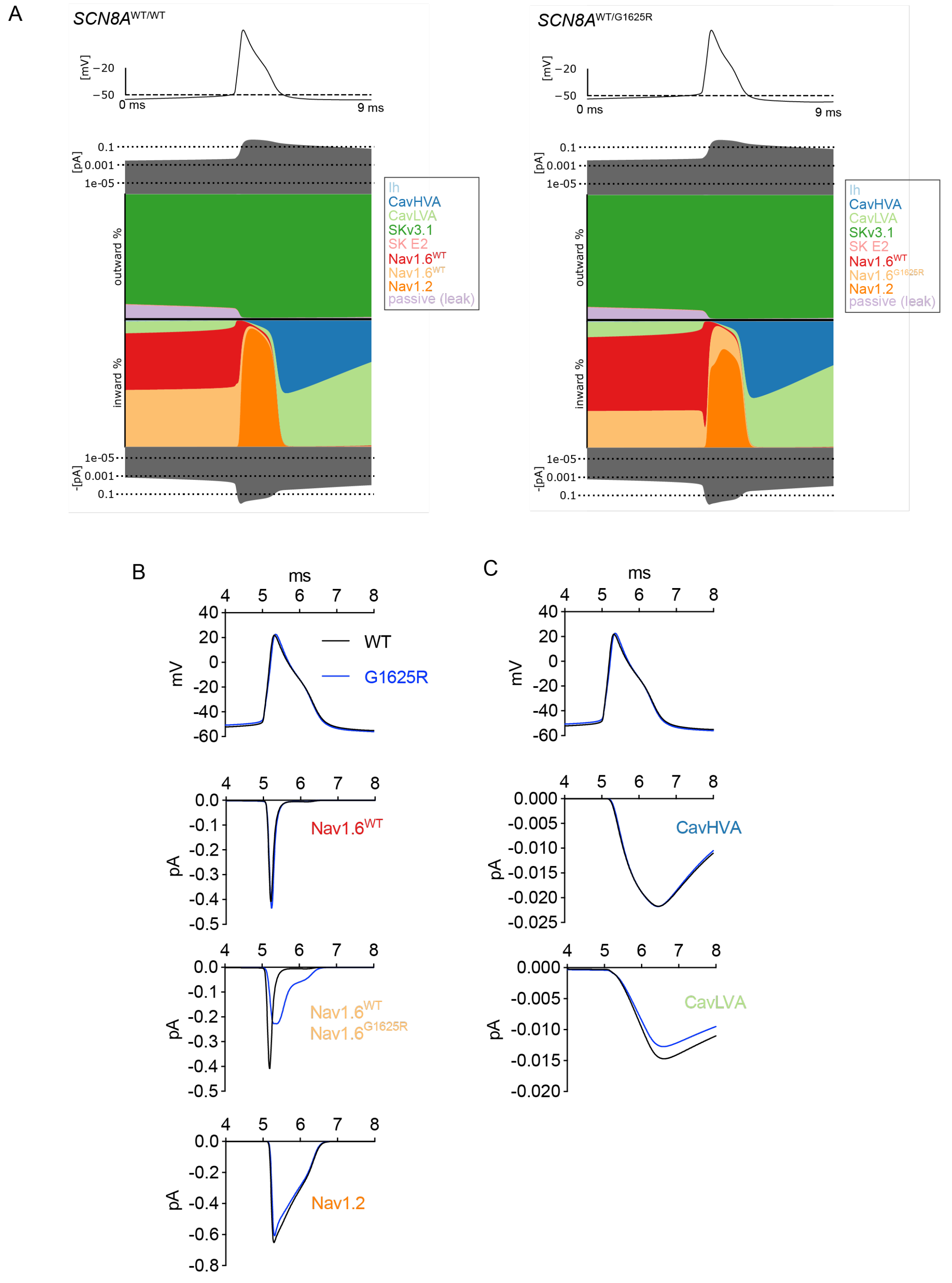


Supplementary Fig. 1. Breakdown of the underlying currents in a cortical pyramidal cell model expressing *SCN8A*$^{WT/WT}$ or *SCN8A*$^{WT/G1625R}$.

**(A)** Breakdown of the ion channels underlying the modeled firing in a patient expressing 100% Nav1.6$^{WT}$ (*SCN8A*$^{WT/WT}$) or 50% Nav1.6$^{WT}$ and 50% Nav1.6$^{G1625R}$ (*SCN8A*$^{WT/G1625R}$). The top trace is the AP; below is the total outward current, the % contribution of each ion channel to the outward and inward current at each time point, and total inward current. Note: Ih has minimal contribution to inward current percentage prior to AP initiation.

**(B)** Modeled sodium currents during the AP. The AP and currents were aligned at the threshold (20 mV/ms).

**(C)** Modeled Calcium currents during the AP. The AP and currents were aligned at the threshold (20 mV/ms).

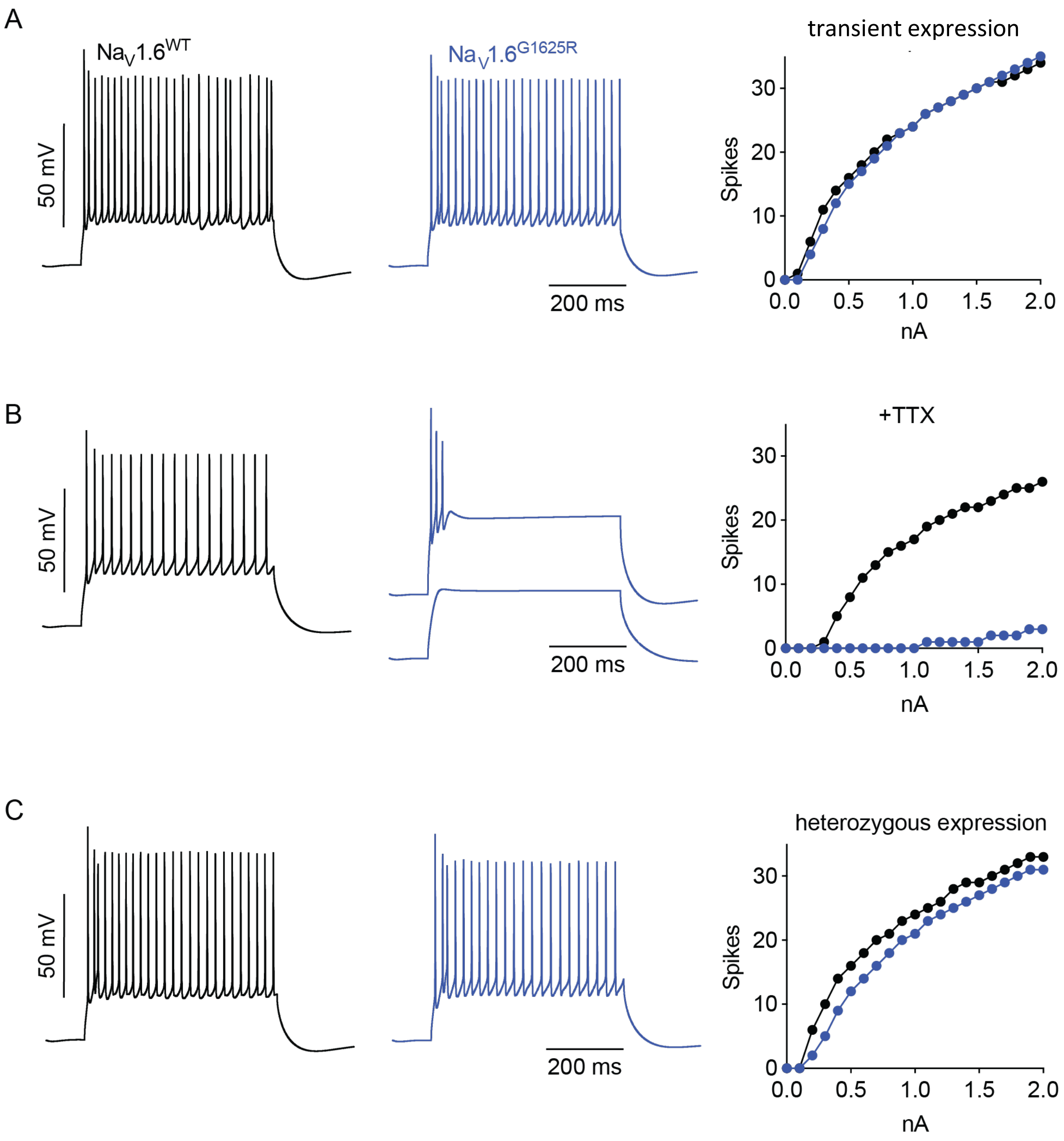


Supplementary Fig. 2. Neuronal excitability in a cortical pyramidal cell model expressing similar current amplitudes of $Na_V1.6^{WT}$ and $Na_V1.6^{G1625R}$. In contrast to the data presented in Fig. 3, in which $Na_V1.6^{G1625R}$ had lower current amplitude, the data in this figure depict the firing in response to neurons expressing similar sodium current amplitudes of both WT and G1625R. **(A)** Modeled firing in response to 20% overexpression of $Na_V1.6^{WT}$ or $Na_V1.6^{G1625R}$, in addition to other endogenous $Na_V$ channels. Examples of modeled firing in response to injection of 1 nA current and F-I curves. **(B)** Modeled firing in response to overexpression of $Na_V1.6^{WT}$ or $Na_V1.6^{G1625R}$, without the contribution of endogenous $Na_V$ channels. Examples of modeled firing in response to injection of 1 nA current (WT and the lower trace in G1625R), or 2 nA (the top trace in G1625R) and F-I curves. **(C)** Modeled firing in a patient expressing 100% $Na_V1.6^{WT}$ or 50% $Na_V1.6^{WT}$ and 50% $Na_V1.6^{G1625R}$. Examples of modeled firing in response to injection of 1 nA current and F-I curves.

### References


Barker, B. S., Ottolini, M., Wagnon, J. L., Hollander, R. M., Meisler, M. H., and Patel, M. K. (2016). The *SCN8A* encephalopathy mutation p.Ile1327Val displays elevated sensitivity to the anticonvulsant phenytoin. *Epilepsia* 57, 1458–1466.

Blanchard, M. G., Willemsen, M. H., Walker, J. B., Dib-Hajj, S. D., Waxman, S. G., Jongmans, M. C. J., et al. (2015). De novo gain-of-function and loss-of-function mutations of *SCN8A* in patients with intellectual disabilities and epilepsy. *J. Med. Genet.* 52, 330–337.

Boßelmann, C. M., Hedrich, U. B. S., Lerche, H., and Pfeifer, N. (2023). Predicting functional effects of ion channel variants using new phenotypic machine learning methods. *PLOS Comput. Biol.* 19, e1010959.

de Kovel, C. G. F., Meisler, M. H., Brilstra, E. H., van Berkestijn, F. M. C., Slot, R. van t., van Lieshout, S., et al. (2014). Characterization of a de novo *SCN8A* mutation in a patient with epileptic encephalopathy. *Epilepsy Res.* 108, 1511–1518..

Estacion, M., O'Brien, J. E., Conravey, A., Hammer, M. F., Waxman, S. G., Dib-Hajj, S. D., et al. (2014). A novel de novo mutation of *SCN8A* (Nav1.6) with enhanced channel activation in a child with epileptic encephalopathy. *Neurobiol. Dis.* 69, 117–123.

Guo, Q. bei, Zhan, L., Xu, H. yan, Gao, Z. bing, and Zheng, Y. ming (2022). *SCN8A* epileptic encephalopathy mutations display a gain-of-function phenotype and divergent sensitivity to antiepileptic drugs. *Acta Pharmacol. Sin.* 43, 3139–3148.

Heyne, H. O., Baez-Nieto, D., Iqbal, S., Palmer, D. S., Brunklaus, A., May, P., et al. (2020). Predicting functional effects of missense variants in voltage-gated sodium and calcium channels. *Sci. Transl. Med.* 12, 6848.

Johannesen, K. M., Liu, Y., Koko, M., Gjerulfsen, C. E., Sonnenberg, L., Schubert, J., et al. (2022). Genotype-phenotype correlations in *SCN8A*-related disorders reveal prognostic and therapeutic implications. *Brain* 145, 2991–3009.

Landrum, M. J., Lee, J. M., Benson, M., Brown, G. R., Chao, C., Chitipiralla, S., et al. (2018). ClinVar: improving access to variant interpretations and supporting evidence. *Nucleic Acids Res.* 46, D1062–D1067.

Lindy, A. S., Stosser, M. B., Butler, E., Downtain-Pickersgill, C., Shanmugham, A., Retterer, K., et al. (2018). Diagnostic outcomes for genetic testing of 70 genes in 8565 patients with epilepsy and neurodevelopmental disorders. *Epilepsia* 59, 1062–1071.

Liu, Y., Koko, M., and Lerche, H. (2021). A *SCN8A* variant associated with severe early onset epilepsy and developmental delay: Loss- or gain-of-function? *Epilepsy Res.* 178, 106824.

Liu, Y., Schubert, J., Sonnenberg, L., Helbig, K. L., Hoei-Hansen, C. E., Koko, M., et al. (2019). Neuronal mechanisms of mutations in *SCN8A* causing epilepsy or intellectual disability. *Brain* 142, 376–390.

Pan, Y., Cummins, T. R., Forsythe, I., and Marra, V. (2020). Distinct functional alterations in *SCN8A* epilepsy mutant channels. *J. Physiol.* 598, 381–401.

Poulin, H., Chahine, M., Toth, K., Young, S., Poulin, H., Chahine, M., et al. (2021). R1617Q epilepsy mutation slows Nav1.6 sodium channel inactivation and increases the persistent current and neuronal firing. *J. Physiol.* 599, 1651–1664.

Reynolds, C., King, M. D., and Gorman, K. M. (2020). The phenotypic spectrum of *SCN2A*-related epilepsy. *Eur. J. Paediatr. Neurol.* 24, 117–122.

Sanders, S. J., Campbell, A. J., Cottrell, J. R., Moller, R. S., Wagner, F. F., Auldridge, A. L., et al. (2018). Progress in understanding and treating *SCN2A*-mediated disorders. *Trends Neurosci.* 41, 442–456.

Sasaki, R., Takano, H., Kamakura, K., Kaida, K., Hirata, A., Saito, M., et al. (1999). A novel mutation in the gene for the adult skeletal muscle sodium channel alpha-subunit (*SCN4A*) that causes paramyotonia congenita of von Eulenburg. *Arch. Neurol.* 56, 692–696.

Solé, L., Wagnon, J. L., and Tamkun, M. M. (2020). Functional analysis of three Nav1.6 mutations causing early infantile epileptic encephalopathy. *Biochim. Biophys. Acta - Mol. Basis Dis.* 1866, 165959.

Tidball, A. M., Lopez-Santiago, L. F., Yuan, Y., Glenn, T. W., Margolis, J. L., Clayton Walker, J., et al. (2020). Variant-specific changes in persistent or resurgent sodium current in *SCN8A*-related epilepsy patient-derived neurons. *Brain* 143, 3025–3040.

Veeramah, K. R., O'Brien, J. E., Meisler, M. H., Cheng, X., Dib-Hajj, S. D., Waxman, S. G., et al. (2012). De novo pathogenic *SCN8A* mutation identified by whole-genome sequencing of a family quartet affected by infantile epileptic encephalopathy and SUDEP. *Am. J. Hum. Genet.* 90, 502–510.

Wagnon, J. L., Barker, B. S., Hounshell, J. A., Haaxma, C. A., Shealy, A., Moss, T., et al. (2016). Pathogenic mechanism of recurrent mutations of *SCN8A* in epileptic encephalopathy. *Ann. Clin. Transl. Neurol.* 3, 114.

Wang, D. W., Crotti, L., Shimizu, W., Pedrazzini, M., Cantu, F., De Filippo, P., et al. (2008). Malignant perinatal variant of long-QT syndrome caused by a profoundly dysfunctional cardiac sodium channel. *Circ. Arrhythm. Electrophysiol.* 1, 370.

Wengert, E. R., Miralles, R. M., Wedgwood, K. C. A., Wagley, P. K., Strohm, S. M., Panchal, P. S., et al. (2021). Somatostatin-positive interneurons contribute to seizures in *SCN8A* epileptic encephalopathy. *J. Neurosci.* 41, 9257–9273.

Wengert, E. R., Saga, A. U., Panchal, P. S., Barker, B. S., and Patel, M. K. (2019). Prax330 reduces persistent and resurgent sodium channel currents and neuronal hyperexcitability of subiculum neurons in a mouse model of *SCN8A* epileptic encephalopathy. *Neuropharmacology*. 158, 107699.

Zaman, T., Abou Tayoun, A., and Goldberg, E. M. (2019). A single-center *SCN8A*-related epilepsy cohort: clinical, genetic, and physiologic characterization. *Ann. Clin. Transl. Neurol.* 6, 1445.
